# Simplified geometric representations of protein structures identify complementary interaction interfaces

**DOI:** 10.1101/2019.12.18.880575

**Authors:** Caitlyn L. McCafferty, Edward M. Marcotte, David W. Taylor

**Affiliations:** Department of Molecular Biosciences, University of Texas at Austin, Austin, TX 78712, USA; Center for Systems and Synthetic Biology, University of Texas at Austin, Austin, TX 78712, USA; Institute for Cellular and Molecular Biology, University of Texas at Austin, Austin, TX 78712, USA; LIVESTRONG Cancer Institutes, Dell Medical School, Austin, TX 78712, USA

## Abstract

Protein-protein interactions are critical to protein function, but three-dimensional (3D) arrangements of interacting proteins have proven hard to predict, even given the identities and 3D structures of the interacting partners. Specifically, identifying the relevant pairwise interaction surfaces remains difficult, often relying on shape complementarity with molecular docking while accounting for molecular motions to optimize rigid 3D translations and rotations. However, such approaches can be computationally expensive, and faster, less accurate approximations may prove useful for large-scale prediction and assembly of 3D structures of multi-protein complexes. We asked if a reduced representation of protein geometry retains enough information about molecular properties to predict pairwise protein interaction interfaces that are tolerant of limited structural rearrangements. Here, we describe a cuboid transformation of 3D protein accessible surfaces on which molecular properties such as charge, hydrophobicity, and mutation rate can be easily mapped, implemented in the MorphProt package. Pairs of surfaces are compared to rapidly assess partner-specific potential surface complementarity. On two available benchmarks of 85 overall known protein complexes, we observed F1 scores (a weighted combination of precision and recall) of 19-34% at correctly identifying protein interaction surfaces, comparable to more computationally intensive 3D docking methods in the annual Critical Assessment of PRedicted Interactions. Furthermore, we examined the effect of molecular motion through normal mode simulation on a benchmark receptor-ligand pair and observed no marked loss of predictive accuracy for distortions of up to 6 Å RMSD. Thus, a cuboid transformation of protein surfaces retains considerable information about surface complementarity, offers enhanced speed of comparison relative to more complex geometric representations, and exhibits tolerance to conformational changes.

## INTRODUCTION

Proteins often assemble into multi-protein complexes as their native forms, mediated by pairwise (or higher-order) protein-protein interactions. Knowledge of the participating protein-protein interfaces involved in forming these complexes is thus critical for understanding and perturbing protein function in a cellular context. Most of our understanding about the contact surfaces by which proteins interact has been from direct experimental determination using techniques such as X-ray crystallography and electron microscopy^1, 2^, but these methods remain costly and laborious. Other, more indirect experimental techniques, including mutagenesis^3, 4^, mass spectrometry^5^, and cross-linking analysis^6^, can also illuminate the specific residues that participate in these interaction interfaces. These techniques give partial information about the three-dimensional (3D) information on the assembly of complexes, and new integrative computational modeling strategies are increasingly able to consider such data as distance restraints to infer 3D structures^7–10^. To complement such experimentally-led approaches, there has also been a strong push to develop better computational approaches for predicting protein interaction interfaces directly from protein amino acid sequences and 3D structures.

Importantly, the prediction of protein-protein interaction interfaces is of substantially lower computational complexity than the problem of predicting or folding a 3D protein structure based on its linear amino acid sequence, as interface predictions (for example, by molecular docking) are limited to 6 degrees of rotational and translation freedom and a sampling of accompanying intramolecular motions that might occur upon binding^11^. Ideally, successful interface predictors would go beyond predicting pairwise interactions and be useful to assemble large molecular machines from individual subunits.

Such predictions are complicated by the fact that protein-protein interactions may take quite different forms, and interactions can be categorized in various ways, including obligate and non-obligate, permanent and transient, and strong and weak^12^. Obligate complexes consist of proteins that are not stable on their own and depend on cooperative folding between the subunits, while non-obligate complexes form from proteins that fold alone and take part in transient or permanent protein interactions. Transient interactions can be further divided into strong and weak interactions. Several studies have determined trends in residues that form protein interfaces. For example, transient interactions have been observed to have similar proportions of hydrophobic residues on both the interaction interface and the remaining surface of the protein. However, because these interfaces are rich in water molecules^13^, there tend to be a larger number of polar residues along the interface^14^. Additionally, many of the forces driving these interactions derive from weak electrostatic charge^15^. Thus, computational approaches face a significant challenge in having to predict contact interfaces that may vary significantly based on the relevant class of protein-protein interaction for any particular interface.

Computational approaches for determining how proteins interact include predictions of interaction interfaces or docking of protein structures, where the former informs the latter. It has been shown that knowledge of an interaction interface can greatly improve the prediction of the conformation of the proteins that are interacting^16^. Interface predictors may be divided into two groups: intrinsic- and template-based approaches^17^. Intrinsic-based approaches focus on features within the protein sequence or the protein structure. Template-based approaches search through databases of protein complexes with known structures and use these interfaces to make predictions^18^. However, the latter approach requires prior structural information for the protein(s) of interest. Intrinsic-based approaches take either sequence information or structural information as the input of the predictor. Enhancing the intrinsic-based approaches may be challenging, as a review of previous literature found that the addition of more features does not improve predictions^17^.

Sequence-based predictors utilize protein sequence information to either feed different amino acid properties into a machine learning classifier or sequence alignment tools. Sequence alignment methods assume that proteins of similar sequences have similar binding partners and therefore binding sites^18^. Many machine learning techniques focus on features of neighboring residues, where the size of the window of residues ranges from 9 to 21 amino acids^18^. However, proximity in sequence does not necessarily reflect proximity in structure, demonstrating one of the benefits of incorporating structural information into the interface predictions. Some techniques have taken an intermediate approach where the proteins are represented by a network where individual nodes represent residues and residue properties, while edges represent structural information providing some spatial resolution^19, 20^.

Structure-based predictors utilize structural information from either experimental data or homology modeling as a constraint in formulating their prediction. Previous studies showed that the quality of the prediction is dependent on the quality of the structure and that homology models produce less accurate predictions^18^. One such structural approach involves dividing a protein surface into patches and using these patches to predict interaction sites. Patches are defined as either the *n* closest residues where *n* depends on the size of the protein or a set size for all proteins^21, 22^. For these methods, patch size is predetermined and uniform, causing problems for predicting interfaces of proteins with multiple binding partners or if the defined surface patch does not accurately reflect the size of the true interface^21^. Many predictors ignore the binding partner; however, utilizing the binding partner has been shown to improve predictions^17^.

Partner-specific interface predictors, which account for all participating proteins in the interaction are less common but have the benefit of considering complementarity between specific proteins. Partner-specific predictors use structures or sequences of two proteins that are assumed to interact in predicting the interaction interface for each protein^17^. A partner-specific approach allows the user to consider complementarity, which plays a central role in molecular recognition. Proteins that promiscuously bind to multiple partners present a unique challenge for predicting interfaces. These multiple binding partners may all bind at the same site, or they may bind at multiple sites on the protein surface^23^. While recent studies highlight the ability of current predictors to separate non-binding from binding residues on individual proteins, these predictors fail to distinguish partner-specific interaction sites resulting in cross-prediction between sites^18^.

Currently, many partner-specific approaches exist for predicting interactions. A majority of these methods use the primary sequence and homology searches to make predictions. PAIRpred utilizes a support vector machine classifier for predicting partner-specific interaction interfaces^24^. While this approach employs multiple features, the features included in the classifier are all based on solvent accessible surface area, which cannot account for proteins that undergo a dramatic conformational change during binding. Another partner-specific tool is PPIPP. PPIPP uses a neural network trained on interacting pairs and has been shown to outperform partner-unaware models^25^. Similarly, HomPPI uses sequence-homology based approaches to identify conserved regions between the partners^26^. Both approaches only use sequence information and do not incorporate spatial data. Many recent approaches have attempted to use multiple sequence alignments (MSAs) to predict residues that coevolve between proteins through direct coupling analysis, mutual information, or a combination of the two and show improved prediction capabilities^8, 27, 28^.

One important challenge that remains for partner-specific, structure-based predictors is accounting for conformational changes that occur upon binding. The performance of these methods decreases with increasing conformational rearrangements and dynamics of the protein pairs upon binding^25^. For this reason, we were interested in developing a reduced representation of protein structural data that does not explicitly consider shape complementarity. Here, we developed and evaluated a protein shape transformation method (MorphProt) that predicts partner-specific interaction interfaces by mapping properties of protein surfaces to cuboids and rapidly testing for complementary surface patches on these reduced geometric representations. MorphProt shows comparable predictive power to a number of more computationally intensive approaches and tolerance to structural rearrangements in the interaction partners.

## MATERIALS AND METHODS

### Benchmark set of protein-protein interactions

To evaluate the quality of the interaction interface predictions from MorphProt, we used a benchmark set of known protein complexes. The benchmark data set for this method was version 5.0 of the widely used protein-protein interaction docking benchmarks^29^. This benchmark set provides a large library of 230 Protein Data Bank^30^ (PDB) files for non-redundant complexes with varying rigidity, as well as enzyme-containing complexes and antibody-antigen complexes. From this set, we extracted 72 complexes for which we were able to obtain mutation rate data (**Supplementary Information**).

In addition to the protein docking benchmark 5.0, we used the protein docking gold standard, the Critical Assessment of PRedicted Interactions (CAPRI) score set^31^. CAPRI provides an expanded benchmark data set for evaluating scoring functions, which includes 15 published CAPRI targets. We analyzed 13 of the 15 targets. The remaining two targets did not have enough sequences to produce reliable mutation rates.

### Calculated properties of surfaces

The properties that were used in these analyses were charge, hydrophobicity, and mutation rate. The atomic charge was calculated using PDB2PQR^32^. PDB2PQR begins by rebuilding missing non-hydrogen atoms using standard amino acid topologies in conjunction with the existing atomic coordinates to determine new positions for the missing atoms. Next, hydrogen atoms are added and positioned to optimize the global hydrogen-bonding network. Finally, PDB2PQR assigns atomic charges and radii based on the AMBER force field. Here, The PDB2PQR program was run using the Opal server.

The Wimley-White hydrophobicity values^33^ were used in determining residue hydrophobicity. These values are semi-empirical and based on the transfer of free energies of polypeptides that show how favorable an amino acid is in a hydrophobic environment. Each atom in the atomic structure was assigned a hydrophobicity value based on the amino acid it was representing.

Finally, the mutation rates were obtained from the ConSurf Database^34^. This database contains information regarding pre-calculated evolutionary conservation scores. The mutation rates stored in the database are calculated using the Rate4Site algorithm^35^. This method evaluates evolutionary mutation rates using a maximum likelihood estimate assuming a stochastic process. Based on this, amino acid replacement probabilities were computed for each branch of the phylogenetic tree. The tree is then used to cluster closely related sequences and find a consensus sequence for each cluster. The consensus sequences are then compared, and each position may be described as variable or conserved. The frequencies are renormalized to represent a number between 1 and 9. Finally, each of the properties described was stored in the surface of the protein structure as part of the appropriate atomic coordinate.

### Protein shape transformation

To reduce the dimensionality of the intricacies of protein shape, we performed a shape transformation of the 3D atomic structure into a cube. To simplify these calculations, we have created a Python library, MorphProt. The input for these calculations is a PDB file (either an atomic structure or homology model), a PQR file, and a conservation file produced by Consurf^34^ when considering mutation rate. First, we extracted the molecular surface using Michel Sanner’s MSMS program^36^, which uses a 1.4 Å diameter sphere to detect the solvent accessible surface area. Next, we calculated a residue depth for all of the amino acids in the protein sequence using the molecular surface. The residue depth was calculated using Biopython^37^ and was evaluated as the average depth of all atoms in a residue from the calculated surface. Amino acids were said to be contributing to the surface of the protein if their residue depth was less than 5 Å from the calculated accessible surface. We extracted the 3D coordinates for all of the atoms that satisfy these surface constraints.

After the atomic coordinates of the surface are extracted, we extract the maximum and minimum for each x, y, and z coordinate as biased centroids, equal to 6. We then used SKLearn^38^ to perform a K-means clustering. We projected each of the clusters onto a 2D surface, creating the face of the cube. Next, we binned each face of the cube into boxes, forming a grid. For these experiments, a 25 Å^2^ box was used, but MorphProt allows for a customizable bin size. For each binned box, we calculated the average of each property that was stored in the box, creating a 2D matrix of values. Here, each matrix represents the face of an unfolded cube and a side of a protein. Finally, each of these numbers in the matrix may be mapped back to a location on the protein surface.

### Protein interaction interface prediction

We computed a 2D cross-correlation, a common pattern recognition and image processing tool, to predict areas of the protein surface with maximum interaction between properties. The cross-correlation was calculated using MorphProt. Because each protein is reduced to a total of 6 matrices, we calculated a total of 36 2D cross-correlations for each pairwise interaction. In addition, we sampled all 10-degree rotations to account, in an approximate fashion, for different orientations or positions of the initial protein structures.

Next, we extracted the top ten maximum interaction scores (high scores) as putative interaction interfaces. The top ten scores represented areas of maximum interaction and complementarity. For properties such as hydrophobicity, we looked for a maximum cross-correlation score as our top score because we are accessing two highly conserved regions that have the same degree of hydrophobicity or a hydrophobic/hydrophilic pocket. For charge, we took the minimum score to represent the charge complementarity that exists between interacting proteins where positively charged surfaces are likely interacting with negatively charged surfaces resulting in a net charge near 0.

After the top ten scores were selected from the cross-correlation matrix, the score was then mapped back to the input matrices to show the position of the matrices that produced the score. Finally, the overlapping position for each matrix is mapped back to the residues in each of the overlapping areas. The final result is a list of residues for each protein that are predicted to be on the partner-specific interaction interface.

### Evaluation of predicted protein interaction interfaces

To evaluate our predictions, we calculated a confusion matrix to classify predicted interface residues as true positives, false positives, false negatives, and true negatives based on the predicted and actual classes. We defined a residue to be on the interaction interface if any atom from the residue is within 10 Å of an atom from the protein it is in complex with. We then evaluated our confusion matrix where the precision, recall, accuracy, and F1 score are defined accordingly:

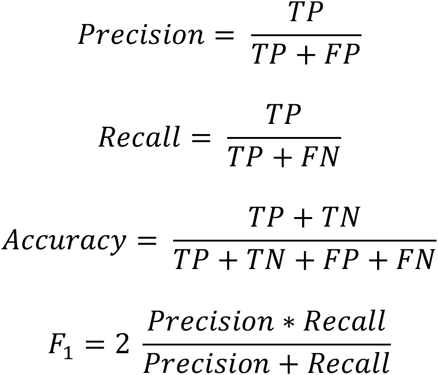

Next, we used an extreme value calculation to validate the “uniqueness” of the atomic properties. We showed that their placement along the interface is not a random distribution of points but rather a clustering of some property. To calculate this, we randomly shuffled the properties associated with each atom and recalculated scores. We repeated this shuffle and scoring 1000 times to generate a distribution. If the score was an extreme value in the distribution, then the score is statistically significant and represented a clustering of a property at that location.

### Simulation of structural distortion by normal mode analysis

To distort the crystal structures from the test set we used elNémo^39^, a normal mode analysis. elNémo predicts the possible movements of a macromolecule through low-frequency normal modes. The l and r unbound subunits of PDBID: 1FQJ from the protein-protein interaction docking benchmark was used. All default parameters were kept except for DQMIN and DQMAX, which were adjusted to 100 and 300, respectively, to allow more extreme distortion. Normal modes 1 and 2 were selected for protein r and normal modes 1 and 4 were selected for protein l. PDBs can be found in the **Supplementary Information**. Modes were selected based on large distortion from RMSD.

## RESULTS

We wished to test if a highly simplified geometric representation of a 3D protein surface embedded with properties was sufficient to predict protein-protein interaction interfaces. The simplification significantly reduces computational complexity, so the question is whether the algorithm would retain its predictive power using the simplified representation and whether the simplified representation would be tolerant of possible molecular motions relevant to the interaction. We wanted to consider protein surface properties and how opposing surfaces complement each other when forming an interface, largely independently of protein shape. For this reason, we began with a transformation of the irregular shape of a protein by considering atoms within 5 Å of the surface of the native protein. This excludes the atoms that play a role in stabilizing the protein core and presumably make less of a contribution to protein-protein interactions.

Our simplified representation is as follows: The solvent accessible surface of the protein is computed and transformed into a simplified geometric representation, the surface of a cuboid, in which the size of the cuboid is proportional to the size of the protein. The transformation thus retains an approximate representation of interface proportions. Recently, the idea of reducing proteins to simplified shapes has gained attention in structural searches^40^. Our shape transformation uses a K-means clustering algorithm to separate protein surface accessible amino acids into 6 distinct clusters, followed by a projection of the coordinates into two-dimensions (2D) (**Fig. 1) to** represent the surfaces. Each atomic coordinate is described by its unique properties. These 2D coordinates are then binned into a grid based on the transformed atomic coordinate locations, and the average property value is calculated for each square of the grid. The result is a matrix of property values where the locations of the values within the matrix represent the neighbors of the atoms on the protein surface with minimal distortion.

**Fig. 1:**
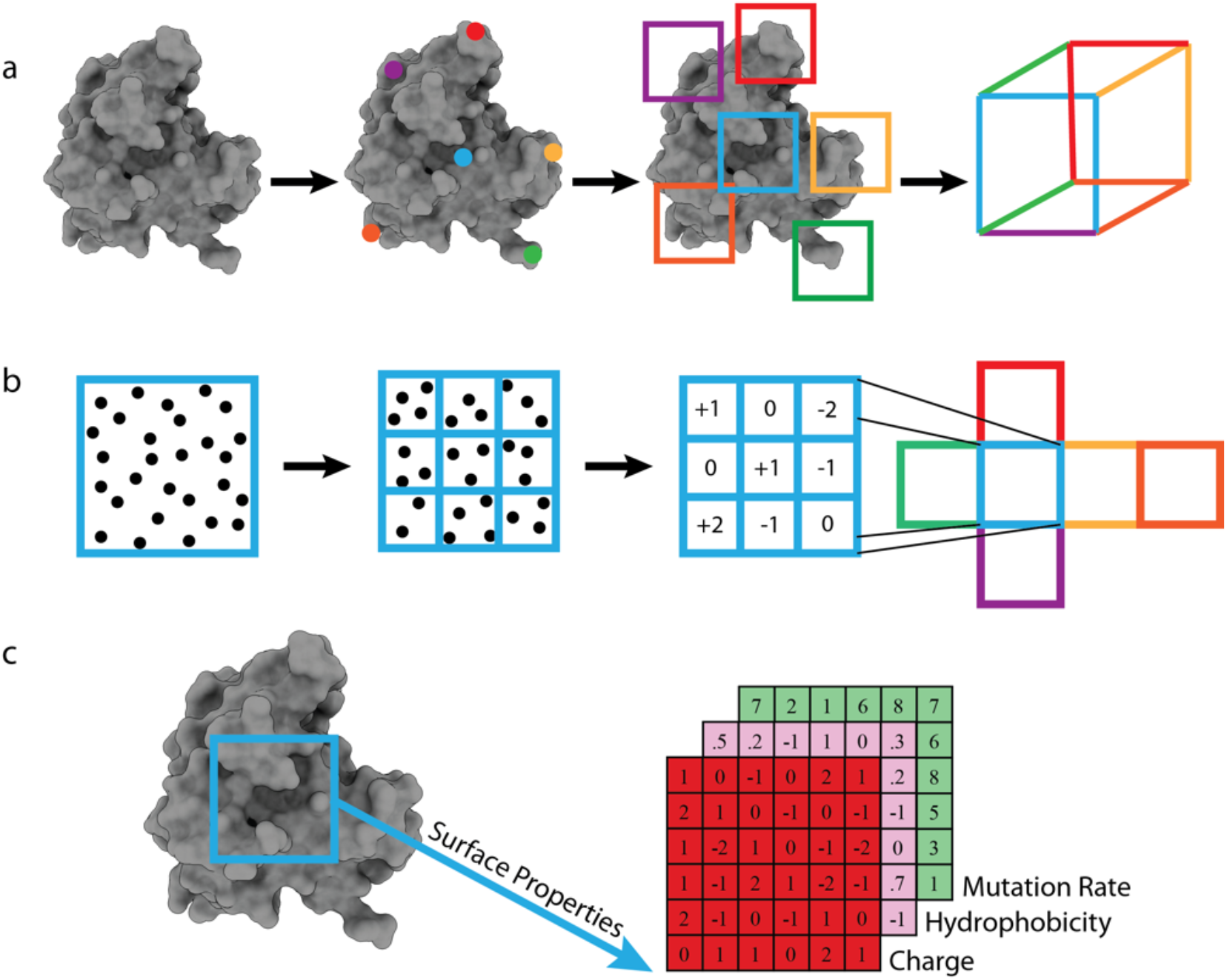
Shape transformation of a protein into a cuboid. **a,** K-means surface clustering of a representative protein structure (PDBID: 1A2K) reduced to a cube. **b,** The atomic properties of each face are binned based on their coordinates (default side length is 5 Å). Average property values are calculated for each box. **c,** The property matrices can be mapped back to the original structure. Potential properties include charge, hydrophobicity, and mutation rate.

These reduced protein surfaces are images, making them suitable for several image processing techniques. To build a partner-specific predictor that considers surface property-complementarity, we performed cross-correlation of images from two partner proteins to find an area of maximum similarity between the two images by computing a dot product at each position after rotation and translation (**Fig. 2**). Cross-correlations have already proven to be invaluable in various image processing techniques, including identifying single particles from electron microscopy data^41^. Here, this approach was used to identify an area of maximum interaction by searching and calculating a complementarity score between properties in the matrix. Because our protein surfaces were reduced into 6 matrices, one representing each side of the cube, we cross-correlated each matrix of one binding partner with each matrix of its partner and generated a score for each position of the 36 cross-correlations. The highest scores represent the positions of each face of the cube where the maximum interaction occurs. The position of the matrices can be mapped back onto the protein surface that they represent. We designed a Python package called MorphProt to perform the shape transformation, cross-correlation evaluations, and plot the predicted interface residues onto the atomic structure.

**Fig. 2:**
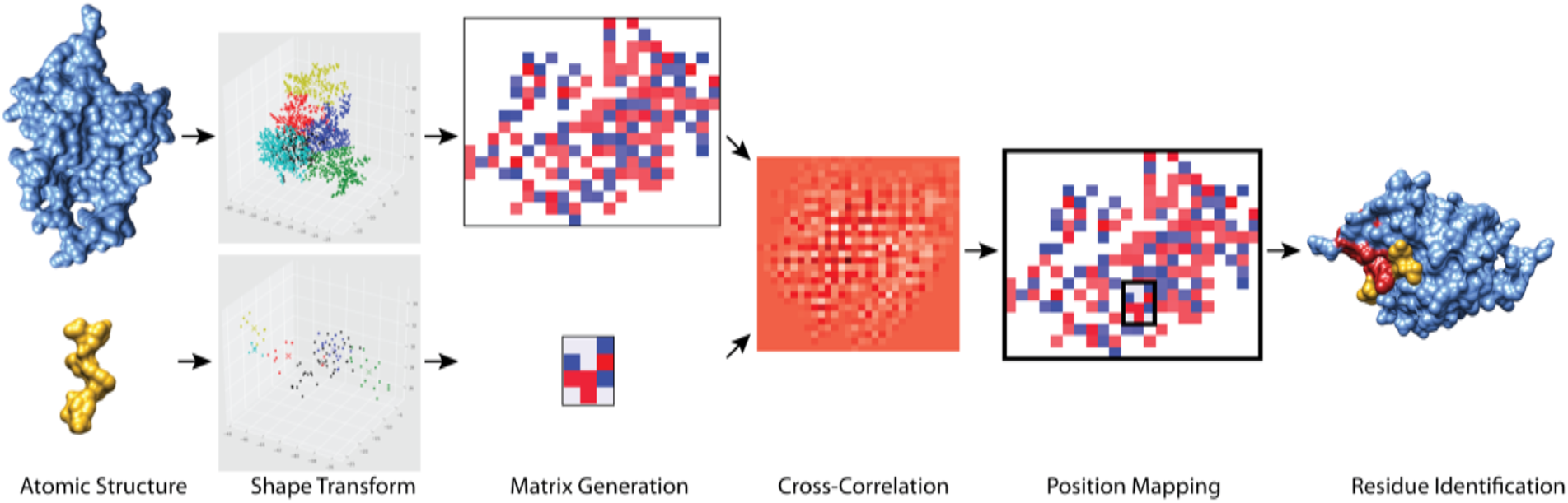
The MorphProt pipeline for interaction interface prediction. The program uses a PDB as input to generate the cube transformation, which is then unwrapped into 6 matrices. Cross-correlations between the faces of the two proteins of interest are calculated. The area of maximum interaction based on complementary is selected from the cross-correlation matrices and mapped back onto the protein structure. In this example, the interaction interface between protein Numb homolog (light blue) and its ligand/inhibitor PTBi peptide (gold) was predicted using charge (PDBID:5NJJ). Positive and negative charges are depicted by red and blue, respectively. In the cross-correlation matrix, the darkest red represents the maximum interaction.

To evaluate the significance of these predictions and their contribution to the protein interface, we used an extreme value approach, which aims to illustrate the distribution of properties across the surface and identify those areas where pockets of each property form. These “property pockets” indicate an area that is likely contributing to a surface interaction. To evaluate this, we randomly shuffled each of the properties to different atomic positions on the protein surface and then recalculated our maximum interaction score with the new distribution of properties. By repeating this process 1000 times, we created a distribution of scores. We selected the unshuffled, predicted high scores from the distribution to determine if it was an extreme value (i.e. in the tail of the distribution). This analysis showed the property of interest is not randomly dispersed across the protein surface; instead, they form pockets, likely occurring on the interaction interface.

To address the concern of any distortion by the shape transformation, we demonstrated that interaction interfaces are still detectable with a proof-of-concept protein pair, the alpha-chymotrypsin-eglin c complex (PDBID:1ACB) (**Fig. 3**). We extracted the surface of each protein in the complex and set the charge property to 0 at all positions with the exception of the true interface. We defined the true interface as all atoms from one protein that are within 10 Å of an atom of the other protein in the complex. The atoms on the true interface of alpha-chymotrypsin were assigned a charge of +1, and those on the true interface of eglin c were assigned a charge of −1. We then performed our shape transformation and cross-correlation analysis using MorphProt. The top ten interaction scores were all between the same two protein faces, which cluster along the true interface. This indicates that despite any distortion that occurs from our reduced representation of the protein surface, MorphProt was still able to identify the area of complementarity between the two surfaces. In addition, when the surface properties were shuffled, the true location of the property was identified as an extreme value. These results further support the notion that the shape transformation does not cause significant distortions and cross-correlation can be used to find the true interface of complementary properties.

**Fig. 3:**
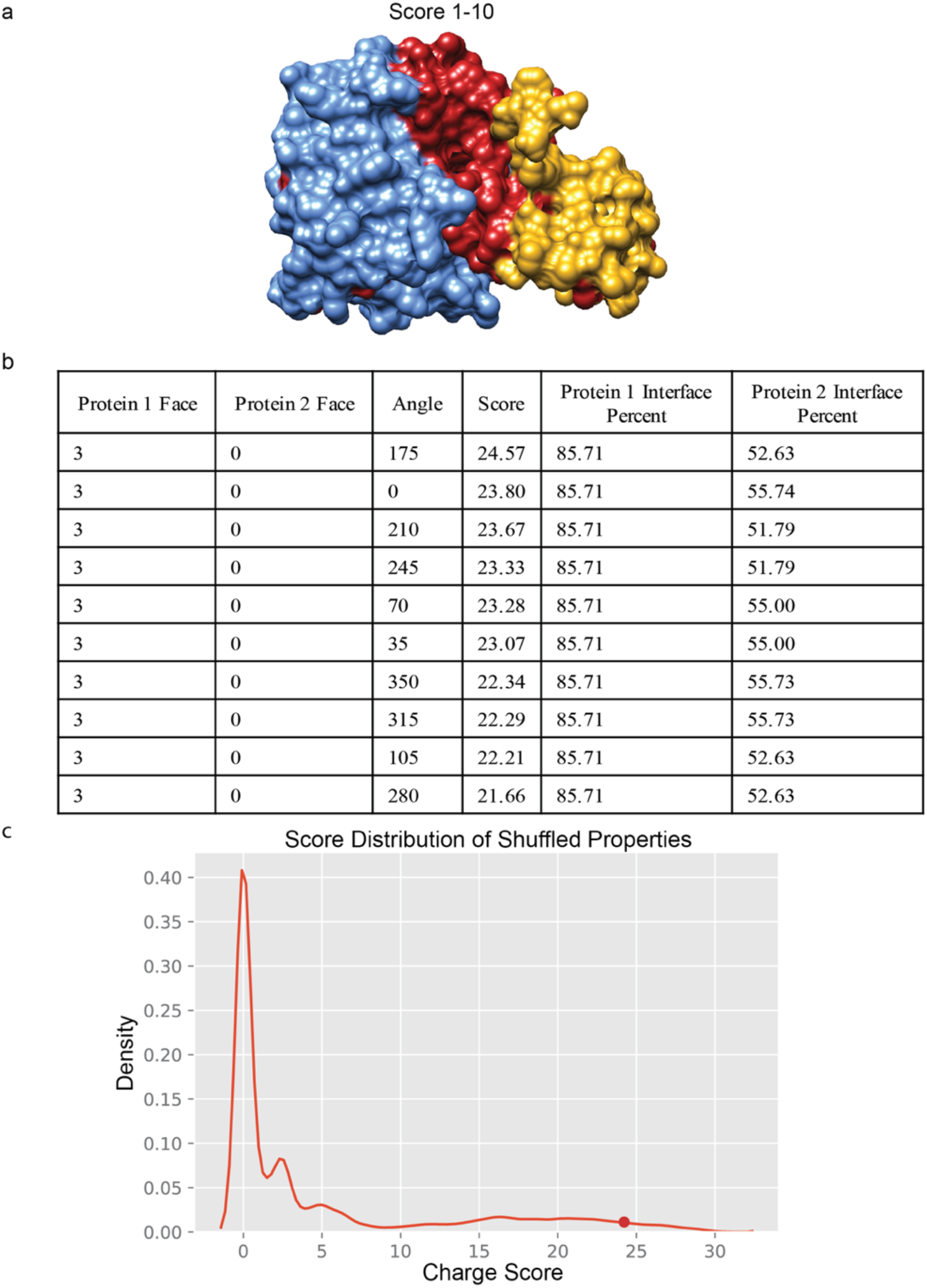
Demonstration of detectable interfaces using the cuboid transformation. **a,** The experimentally determined structure of the alpha-chymotrypsin-elgin c protein complex (PDBID:1ACB). All charge values were set to 0 on the surface of both proteins. The ligand or **l** (gold) interface residues were set to −1 and the receptor or **r** (light blue) interface residues were set to +1. The predicted interface (red) was mapped onto the protein complex. **b,** The table shows the faces of the top 10 scores. The interface percent shows the percent of residues that are within 10 Å of the partner protein. **c,** The cross-correlation scores produced from 1000 shuffles of the engineered charge property across the surface of the protein. The point represents the top score from the prediction.

Next, our partner-specific interaction interface predictor was used to predict the interfaces of the CAPRI score set^31^, a gold standard in protein docking. We predicted the interaction interface according to charge, hydrophobicity, and mutation rate of the unbound structures and mapped the prediction onto the interface of the bound structures (**Fig. 4b**). The interaction interface predictions were scored based on the number of true positives, precision, accuracy, and F1score for the top ten scores. The true positive, false negative, and false positive predictions are defined in **Fig. 4a** for each predicted interface (see **Methods**). The number of true positives reflects the sum of all correct predictions in the dataset. The precision, accuracy, and F1 score represent the average across the CAPRI dataset. The individual CAPRI statistics were also calculated (**Supplementary Information**). Overall, mutation rate is the most predictive property based on surface complementarity with an average accuracy of 61% and F1 of 28%. For charge, hydrophobicity, and mutation rate the average precision was 35%, 33%, and 42% and the average F1 score was 21%, 19%, and 28%, respectively. However, on a case-by-case basis, different properties can provide the best prediction for certain complexes. For example, in the prediction of the interface of the colicin-E2 immunity protein and the colicin-E9 complex (PDBID: 2WPT, Target ID: T41), charge and hydrophobicity prove to be the most predictive properties with accuracies and F1 scores 10% higher than the predictions from mutation rate. Further examination of this complex shows that the complex is non-cognate, which explains why mutation rate is a poor predictor. Additionally, there is a disulfide bond and extensive hydrogen bonding between the interface of the two proteins^42^, hence the improved prediction quality of the charge and hydrophobicity based properties. In addition to the CAPRI score set, we evaluated this approach on 72 of the integrated protein-protein interaction benchmark complexes (**Supplementary Information**)^29^. We obtained similar results to the CAPRI data set for the protein-protein interaction benchmark where the average precision was 35%, 31%, and 48%, and F1 score was 23%, 21%, and 34% for charge, hydrophobicity, and mutation rate, respectively. However, individual property predictions displayed precision and F1 scores as high as 86% and 56% for mutation rate, 74% and 39% for charge, and 67% and 48% for hydrophobicity. Taken together, MorphProt can predict interaction interfaces based on surface property complementarity despite a loss of structural information.

**Fig. 4:**
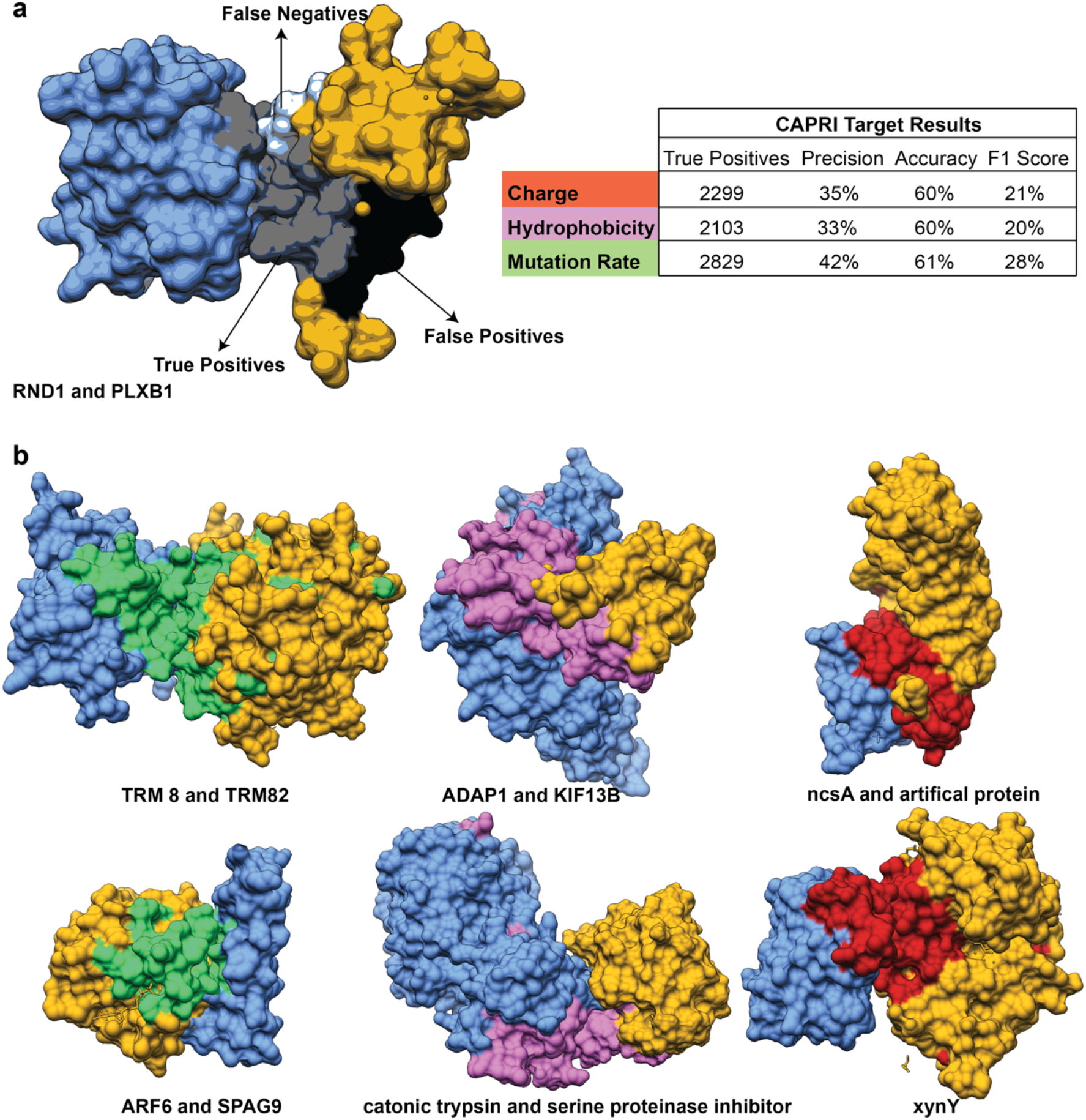
MorphProt validation with the CAPRI score set. **a,** True positive, false negative, and false positive residue predictions are shown for a representative protein (PDBID: 2REX). The results for the CAPRI score set are summarized in the table. Red, pink, and green represent charge, hydrophobicity, and mutation rate, respectively. **b,** Results of six representative CAPRI score set protein complexes (PDBID: 2VDU, 3FM8, 4JW3, 2W83, 3E8L, 2W5F) are depicted.

Of primary interest for biological processes, is the assembly of large macromolecular complexes. Using Morphprot, we can perform pairwise predictions with knowledge of subunits that are directly interacting by indirect methods. We explored the assembly of a large protein complex by examining our recently published Ceru+32/GFP-17 protomer structure^43^, a synthetically engineered supercharged GFP 16-mer. These proteins were engineered to have oppositely charged variants of the normally monomeric green fluorescent proteins (GFP), which resulted in the assembly of a large, ordered macromolecular structure. Because the structure is known to form charge-based interactions, it served as an effective test for the ability of MorphProt to predict partner-specific interactions within a large macromolecular complex where subunits have multiple interaction interfaces. The input for MorphProt was the α and β supercharged subunits. The top ten scores accurately predicted both of the charge-based interfaces between subunits (**Fig. 5**).

**Fig. 5:**
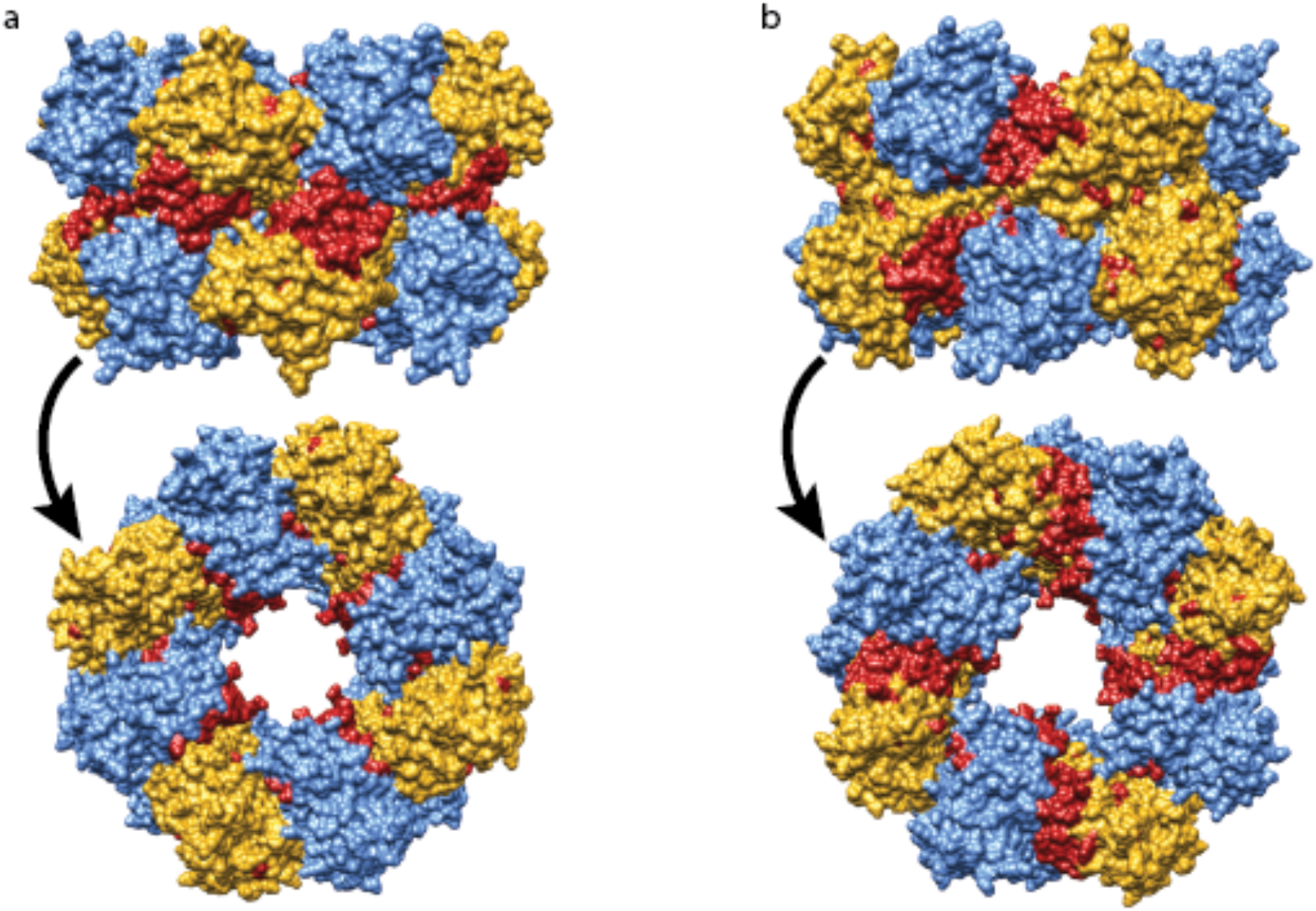
An example of MorphProt applied to predict contact interfaces in a large multimeric protein assembly. MorphProt predicts interfaces of the Ceru+32/GFP-17 protomer (PDBID: 6MDR) between the alpha and beta subunits. The charge predicted interface is shown (red) for **a** score 2 and **b** score 9 predictions of the top 10 scores.

To demonstrate the advantages of using a partner-specific, surface property complementarity method, we considered two binding scenarios that present challenges for conventional interface predictors: (1) a protein that has multiple binding partners and sites and (2) a protein that undergoes a dramatic conformational change upon binding to a partner. To test the multiple-binding site scenario, we used the lysozyme and anti-lysozyme complex (PDBID: 1BVK). The heavy and light chains of the anti-lysozyme form a hydrophobic zipper upon cooperative folding ^44^ and interact with their antigen, lysozyme (**Fig. 6**). Here, we accurately predicted the hydrophobic interaction between the heavy and light chains of the antibody and the charge-driven interaction between the antibody and antigen. To validate that our algorithm can handle dramatic structural rearrangements, we tested the interleukin-1 receptor and the interleukin-1 receptor antagonist complex (PDBID: 1IRA), where the interleukin-1 receptor undergoes a dramatic conformational change upon complex formation (approx. 26.2 Å across all residue pairs). Again, we were able to accurately predict the interaction interface between the protein pair despite this large-scale structural rearrangement.

**Fig. 6:**
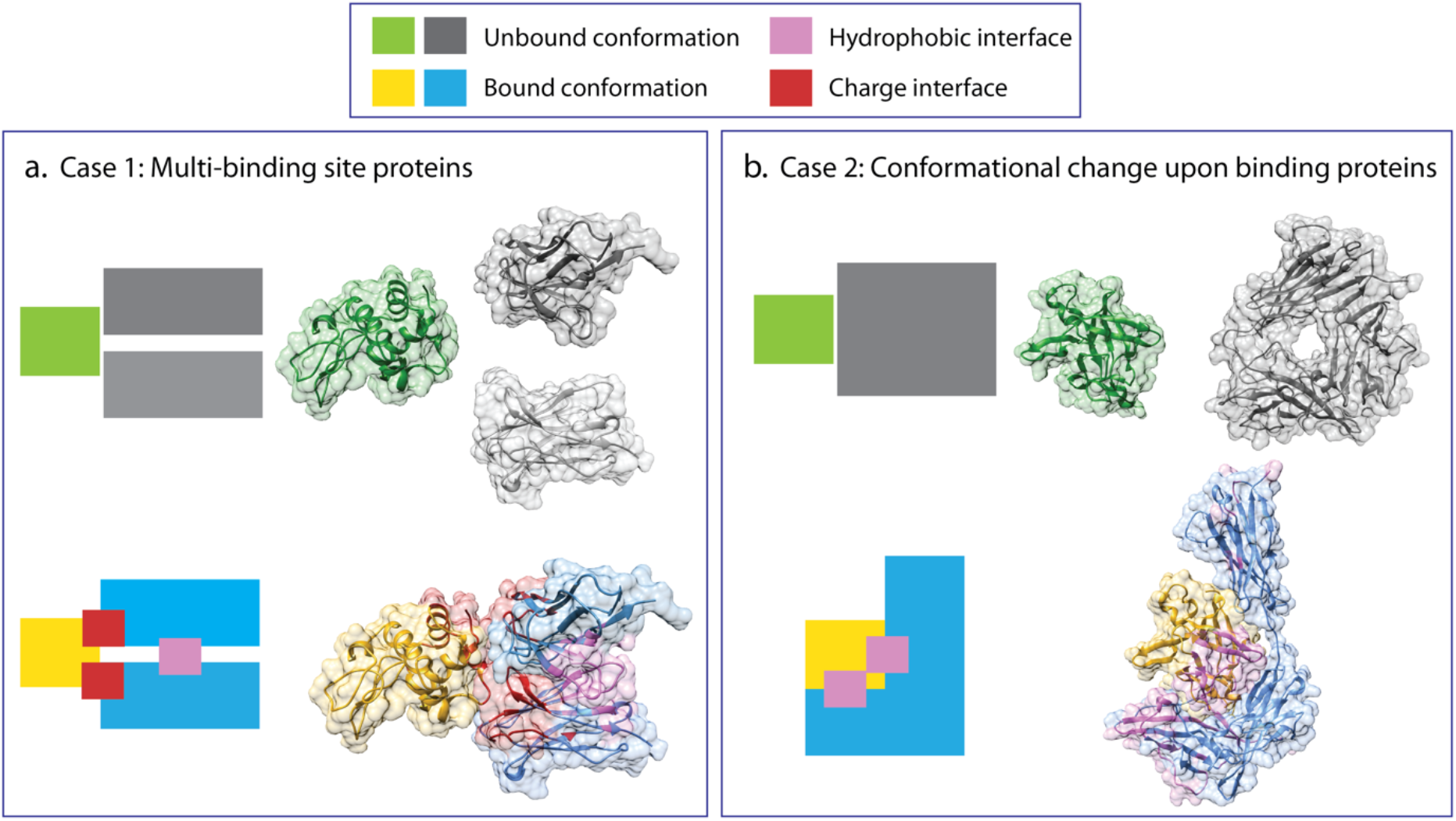
MorphProt successfully predicts interfaces for challenging binding scenarios. **a,** For proteins with multiple binding sites, MorphProt can predict each distinct partner-specific interaction interface. Shown is the antibody-antigen interaction between lysozyme and anti-lysozyme (PDBID: 1BVK). The interaction interface between the heavy chain and the light chain of the anti-lysozyme is predicted using hydrophobicity, while the interaction interface between the anti-lysozyme chains and lysozyme are predicted using charge. **b,** Because MorphProt utilizes charge, hydrophobicity, and mutation rate to predict interfaces, it accurately predicts binding pockets for proteins that undergo dramatic structural rearrangements. Depicted is the interleukin-1 receptor and the interleukin-1 receptor–antagonist complex (PDBID:1IRA) interface. As shown from the unbound and bound structure, the receptor undergoes a striking conformational change upon antagonist binding.

Finally, we wanted to test the performance of our interface predictor on uncertain structural models produced by homology modeling or other structural prediction algorithms. In both experimental and computational structure building, there can occasionally be uncertainty regarding the exact position of the side chains and backbone of the model. By distorting one of our test proteins that produced a strong mutation rate interface prediction, we showed that our predictions remain robust even considering a structure that is distorted by up to ~6 Å (**Fig. 7**). The crystal structures of the unbound Gnai and RGS9 (PDBID: 1FQJ) were distorted using normal mode analysis. We used elNémo^39^ to compute the low-frequency normal modes of each of the structures in the complex. In the analysis, one of the subunits (receptor or ligand) was held constant, while the interface was predicted at different RMSD distortions of the other subunit (receptor or ligand). Despite different configurations of the protein backbone, we were still able to predict the interface based on the generalized property complementarity for a given section of the protein structure.

**Fig. 7:**
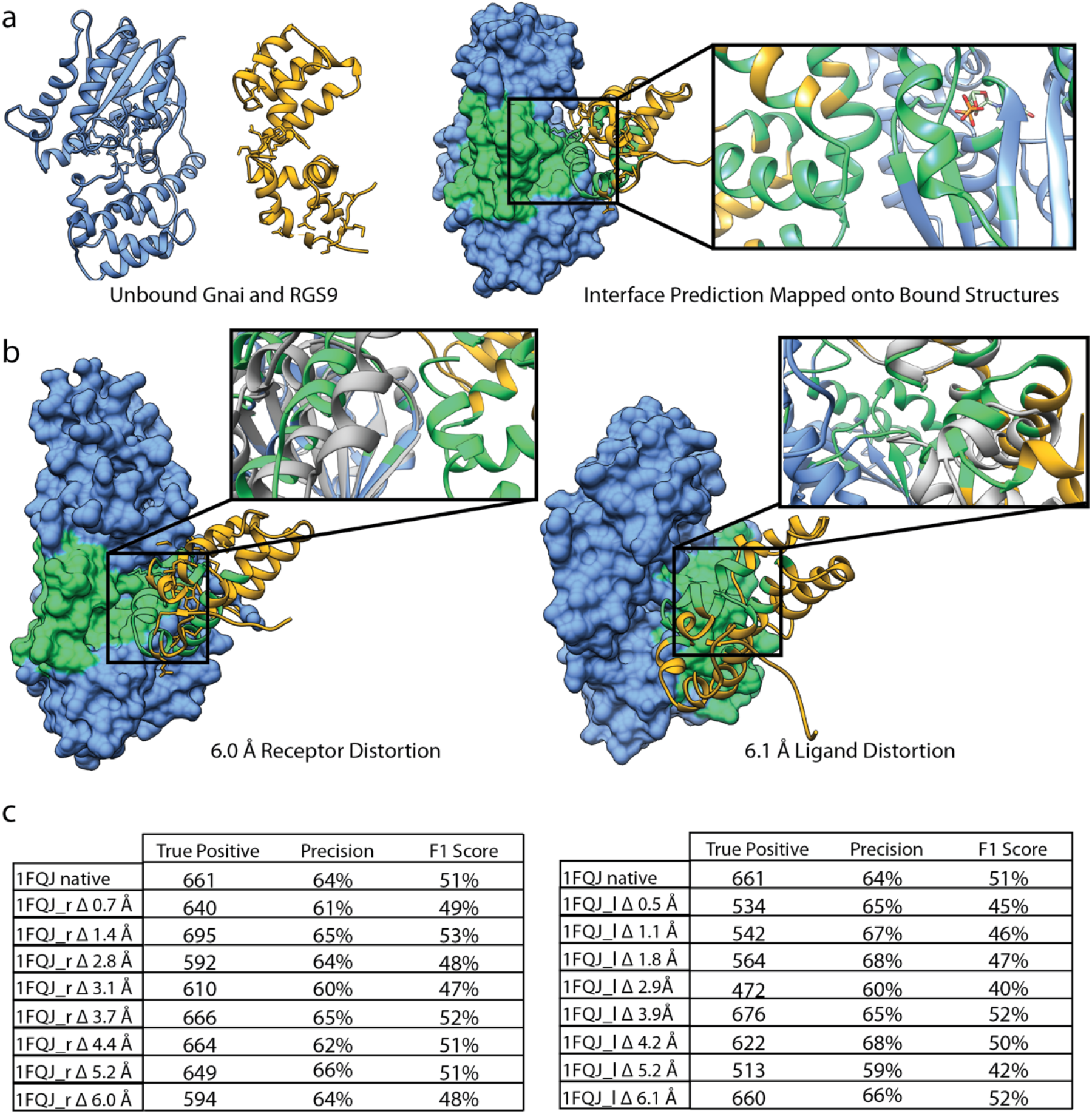
MorphProt can predict interaction interfaces despite structural distortion. **a,** Unbound structure of Gnai and RGS9 (PDBID: 1FQJ). The ligand and receptor are depicted in gold and blue, respectively. The interface is predicted using mutation rate. The predicted interface is colored green on the bound structure. **b,** The receptor and ligand were distorted using elNémo normal mode analysis. While the receptor was distorted up to ~6 Å, the ligand was held constant and vice versa. The interfaces were again predicted using mutation rate and the distorted structure. The close-up depicts the native structure (grey) superimposed onto the distorted structure to show the change in position of residues on the interface. The predicted interface is mapped onto the distorted structure. **c,** The precision-recall curves and diagnostic table show that there is little change in the prediction despite structural distortion for multiple distorted protein structures. The true positive, false, positive, and false negative parameters are illustrated in **Fig.3**.

## DISCUSSION

Here, we have demonstrated that by using a cuboid transformation to normalize the highly variable 3D protein structure to a simplified geometric shape, we are able to store layers of information on a 2D representation of a protein surface while preserving atomic neighborhoods. The resulting matrix of values contains the location of surface properties and their proximity to other values and is a direct representation of the spatial coordinates of the 3D atomic structure. We showed that converting the surface properties to an image allows us to identify areas of maximum interaction of surface properties between two proteins via a partner-specific approach. We showed that MorphProt can also be used to construct large macromolecular assemblies.

While primary sequences provide information regarding amino acid identity and adjacent residues, it can be difficult to precisely determine from sequence alone which residues reside on the surface of a protein and their relation to each other in its 3D structure. Structure-based approaches allow us to extract and investigate surface properties, providing a useful first step for interface prediction, as the spatial position of residues is essential for macromolecular recognition^22^. Many machine learning interaction interface predictors exist and use structure, but the only information stored in feature vectors is statistical information for the surface patches and not the spatial arrangement of the residues^22^. In addition to the lack of information regarding residue neighborhoods, many of the structure-based approaches are not equipped to handle dramatic conformational changes upon binding^45^. We have addressed these limitations of previous methods through our shape transform by treating the protein surface as a simple 2D matrix, where the location of a value within the matrix is a representation of the location of that value on the protein surface. This novel surface-patch approach turns out to be incredibly powerful in identifying the areas of maximum interaction between structures of interacting pairs.

In our approach, patch size is not predetermined; instead, it is dependent on the size of the proteins being tested and the size of overlap between protein faces for each score calculation. Traditional approaches for identifying a surface patch result in fairly uniform patch sizes^21^. Our method tests surface patches over a number of different sizes and arrangements because the patches are determined by the position of the cross-correlation. The first patch tested is the corner of two matrices and expands as the calculation continues, and the patches are both rotated and adjusted in size. The result is a sample of various patches and orientations, which can be used to identify the area of maximum interaction between the pair of proteins.

In most structure-based interaction interface predictors, an interface is identified based on features of a given area on one of the protein surfaces, ignoring properties of a partner when determining how they best fit together. A partner-specific predictor uses information regarding both proteins of interest. It has been shown that prediction methods that do not employ a partner-specific approach have lower reliability in predicting transient binding sites^46, 47^, whereas a partner-specific approach can identify locations that are highly conserved for transient protein-protein interactions^26^. A significant advantage of using a partner-specific predictor is its ability to find specific surface areas that form interactions with different partners. One significant challenge of many partner-specific predictors is their use of unbound protein structures to search for interaction interfaces^17^. In many biological processes, proteins undergo a dramatic conformational change upon binding, which complicates predicting an interface based on unbound structures. We have demonstrated that using a reduced surface representation of a protein in combination with stored information of highly predictive properties, we can make partner-specific interface predictions for unbound proteins, including those that undergo at least moderate structural rearrangements, an important feature for building multi-protein assemblies.

Our results using MorphProt are promising when compared to other available partner-specific interaction predictors. The developers of PAIRpred reported the identification of a true positive in the top 15 predictions for 7 of the 9 complexes tested using the CAPRI score-set. PAIRpred was unable to predict for targets 3FM8 (T39) and 2VDU (T29). These targets have been reported as being challenging complexes to evaluate in CAPRI^24^. However, MorphProt yielded accurate predictions for 2VDU based on mutation rate (78% accuracy and F1 score of 48%) and 3FM8 based on hydrophobicity (73% accuracy and F1 score of 25%). From the results of CAPRI rounds 15-19, T32, T35, T36, T38, and T39 presented the greatest challenges for docking predictors^48^. While we did not test T35 or T38, our interface predictions for the remaining targets remain robust. 3BX1 had an interaction interface prediction accuracy of 65% and F1 score of 24%, while 2W5F (T36) had an interaction interface prediction accuracy of 73% and an F1 score of 21% (T39 summarized above). Taken together, these results show that MorphProt can perform accurate and precise interface predictions for some of the most challenging CAPRI targets despite not considering shape.

Furthermore, we showed that despite introduced structural distortion, we are still able to predict interfaces based on complementarity. This is increasingly important for predicting interaction interfaces with the widespread use of homology models and lower-resolution structures. Here, greater weight is put on the neighborhoods of properties on the surface rather than their exact location. The ability to predict the interface for homology models is significant for assembling macromolecular complexes where little is known regarding the structure of the individual subunits. Theoretically, one could produce models for the subunits and then arrange them according to their interaction interfaces to predict the structure for large assemblies. Such analyses would also benefit from protein docking following the interface prediction to improve positioning.

While discrepancies between interface prediction and protein docking occur often, the techniques are effective when used in conjunction with one another. Protein-protein docking is a partner-specific technique that is highly dependent on shape complementarity and energetics^22^. One of the limitations of protein-protein docking is the large sample size that must be tested and then scored by an energy function to produce a prediction of the arrangement of two proteins. The number of arrangements would be drastically reduced by using an interface prediction as a preliminary step before docking. Previous studies showed that using a partner-specific, homology-based interface prediction prior to docking significantly improves the scoring of the docked proteins^49^. Notably, the HADDOCK server allows for the incorporation of a predicted interaction interface, however, this interface is computed from a single protein rather than a partner-specific interface^50^. Incorporating our interface prediction into a protein-protein docking pipeline would increase computational efficiency because it is independent of shape complementarity and energetics.

Another significant application of partner-specific interaction interface predictor is the screening of small molecule inhibitors or drugs. These often interact via transient interactions^22^, making predicting transient interactions imperative. Small molecules that interact with protein-protein interfaces and alter these interactions have demonstrated to be effective drugs and the prediction of these interfaces could be useful in finding potential targets^51^. This poses a challenge because many protein interfaces have been described as large, featureless surfaces that lack obvious binding pockets^52^. Because our method reduces the protein surface to essentially the same, we would likely be able to make more accurate predictions using physicochemical properties stored on the surface of the protein. Furthermore, predictions and scores for small molecule inhibitors or drugs could be optimized by understanding the area of interaction produced by our method.

## CONCLUSIONS

To address the inherent variability of protein shape, conformational changes, and structural approximations while reducing computation time, we were interested in determining if a simplified geometry retains enough spatial information to predict interaction interfaces based on complementary properties. Specifically, our aim was to develop a pipeline that was robust to molecular motions while gaining computational power to assemble larger multimeric protein complexes. Using MorphProt, we performed a cuboid transformation of the accessible surface of a protein into the surface of a cuboid. This reduced representation allows for easy storage of intrinsic properties of the protein such as hydrophobicity, charge, and mutation rate to be embedded within each surface image. The result is a quantitative description of these properties across a protein surface enabling image processing techniques to identify complementarity between the properties of two interacting protein surfaces. We show this method can be useful when one of the above properties is the driving force of the interaction. MorphProt was able to predict interaction interfaces for the unbound CAPRI targets and the protein-protein interaction benchmark complexes with comparable results to a number of other predictors. Additionally, MorphProt was able to predict interfaces for a large 16-subunit oligomer, proteins with multiple binding sites, and crystal structures that have been distorted by up to ~6 Å to mimic models built from lower resolution density maps or imperfect homology models, demonstrating a utility to integrated platforms that aim to assemble complicated protein complexes.

## Supporting information

Confusion matrix results for CAPRI and protein docking benchmark 5.0 sets.

PDB files for normal mode analysis of PDBID: 1FQJ.

PDB, PQR, mutation rate, and confusion matrix files of the protein docking benchmark 5.0.

## ACKNOWLEDGEMENTS

The MorphProt package is available at https://github.com/cmccaffe/MorphProt. This work was supported in part by Welch Foundation Research Grants F-1938 (to D.W.T.) and F-1515 (to E.M.M.), National Institutes of Health (R35 GM122480, R01 HD085901, and R01 DK110520 to E. M. M.), Army Research Office Grant W911NF-15-1-0120 (to D.W.T.), and a Robert J. Kleberg, Jr. and Helen C. Kleberg Foundation Medical Research Award (to D.W.T.). C.L.M is an NSF Graduate Research Fellow supported by the National Science Foundation (2019238253), D.W.T is a CPRIT Scholar supported by the Cancer Prevention and Research Institute of Texas (RR160088) and an Army Young Investigator supported by the Army Research Office (W911NF-19-1-0021).

## REFERENCES

1. Bai, X.-c.; Fernandez, I. S.; McMullan, G.; Scheres, S. H., Ribosome structures to near-atomic resolution from thirty thousand cryo-EM particles. elife 2013, 2, e00461.

2. Lowe, J.; Stock, D.; Jap, B.; Zwickl, P.; Baumeister, W.; Huber, R., Crystal structure of the 20S proteasome from the archaeon T. acidophilum at 3.4 A resolution. Science 1995, 268, 533–539.

3. Kortemme, T.; Kim, D. E.; Baker, D., Computational alanine scanning of protein-protein interfaces. Sci. STKE 2004, 2004, pl2–pl2.

4. Schreiber, G.; Fersht, A. R., Energetics of protein-protein interactions: Analysis ofthe Barnase-Barstar interface by single mutations and double mutant cycles. Journal of molecular biology 1995, 248, 478–486.

5. Wan, C.; Borgeson, B.; Phanse, S.; Tu, F.; Drew, K.; Clark, G.; Xiong, X.; Kagan, O.; Kwan, J.; Bezginov, A., Panorama of ancient metazoan macromolecular complexes. Nature 2015, 525, 339.

6. Jessulat, M.; Pitre, S.; Gui, Y.; Hooshyar, M.; Omidi, K.; Samanfar, B.; Tan, L. H.; Alamgir, M.; Green, J.; Dehne, F., Recent advances in protein–protein interaction prediction: experimental and computational methods. Expert opinion on drug discovery 2011, 6, 921–935.

7. Russel, D.; Lasker, K.; Webb, B.; Velázquez-Muriel, J.; Tjioe, E.; Schneidman-Duhovny, D.; Peterson, B.; Sali, A., Putting the pieces together: integrative modeling platform software for structure determination of macromolecular assemblies. PLoS biology 2012, 10, e1001244.

8. Cong, Q.; Anishchenko, I.; Ovchinnikov, S.; Baker, D., Protein interaction networks revealed by proteome coevolution. Science 2019, 365, 185–189.

9. Leitner, A.; Faini, M.; Stengel, F.; Aebersold, R., Crosslinking and mass spectrometry: an integrated technology to understand the structure and function of molecular machines. Trends in biochemical sciences 2016, 41, 20–32.

10. Braitbard, M.; Schneidman-Duhovny, D.; Kalisman, N., Integrative structure modeling: overview and assessment. Annual review of biochemistry 2019, 88.

11. Xu, D.; Tsai, C.-J.; Nussinov, R., Hydrogen bonds and salt bridges across protein-protein interfaces. Protein engineering 1997, 10, 999–1012.

12. Acuner Ozbabacan, S. E.; Engin, H. B.; Gursoy, A.; Keskin, O., Transient protein– protein interactions. Protein engineering, design and selection 2011, 24, 635–648.

13. Perkins, J. R.; Diboun, I.; Dessailly, B. H.; Lees, J. G.; Orengo, C., Transient protein-protein interactions: structural, functional, and network properties. Structure 2010, 18, 1233–1243.

14. Kuroda, D.; Gray, J. J., Shape complementarity and hydrogen bond preferences in protein–protein interfaces: implications for antibody modeling and protein–protein docking. Bioinformatics 2016, 32, 2451–2456.

15. Tang, C.; Iwahara, J.; Clore, G. M., Visualization of transient encounter complexes in protein–protein association. Nature 2006, 444, 383.

16. Xue, L. C.; Jordan, R. A.; El-Manzalawy, Y.; Dobbs, D.; Honavar, V. Ranking docked models of protein-protein complexes using predicted partner-specific protein-protein interfaces: a preliminary study. In Proceedings of the 2nd ACM Conference on Bioinformatics, Computational Biology and Biomedicine, 2011; ACM: 2011; pp 441–445.

17. Esmaielbeiki, R.; Krawczyk, K.; Knapp, B.; Nebel, J.-C.; Deane, C. M., Progress and challenges in predicting protein interfaces. Briefings in bioinformatics 2015, 17, 117–131.

18. Zhang, J.; Kurgan, L., Review and comparative assessment of sequence-based predictors of protein-binding residues. Briefings in bioinformatics 2017, 19, 821–837.

19. Hamer, R.; Luo, Q.; Armitage, J. P.; Reinert, G.; Deane, C. M., i-Patch: Interprotein contact prediction using local network information. Proteins: Structure, Function, and Bioinformatics 2010, 78, 2781–2797.

20. Fout, A.; Byrd, J.; Shariat, B.; Ben-Hur, A. Protein interface prediction using graph convolutional networks. In Advances in Neural Information Processing Systems, 2017; 2017; pp 6530–6539.

21. Jones, S.; Thornton, J. M., Prediction of protein-protein interaction sites using patch analysis. Journal of molecular biology 1997, 272, 133–143.

22. Xue, L. C.; Dobbs, D.; Bonvin, A. M.; Honavar, V., Computational prediction of protein interfaces: A review of data driven methods. FEBS letters 2015, 589, 3516–3526.

23. Teppa, E.; Zea, D. J.; Marino-Buslje, C., Protein–protein interactions leave evolutionary footprints: High molecular coevolution at the core of interfaces. Protein Science 2017, 26, 2438–2444.

24. Afsar Minhas, F. u.A.; Geiss, B. J.; Ben-Hur, A., PAIRpred: Partner-specific prediction of interacting residues from sequence and structure. Proteins: Structure, Function, and Bioinformatics 2014, 82, 1142–1155.

25. Ahmad, S.; Mizuguchi, K., Partner-aware prediction of interacting residues in protein-protein complexes from sequence data. PLoS One 2011, 6, e29104.

26. Xue, L. C.; Dobbs, D.; Honavar, V., HomPPI: a class of sequence homology based protein-protein interface prediction methods. BMC bioinformatics 2011, 12, 244.

27. Mishra, S. K.; Cooper, S. J.; Parks, J. M.; Mitchell, J. C., Hotspot coevolution at protein-protein interfaces is a key identifier of native protein complexes. BioRxiv 2019, 698233.

28. Vajdi, A.; Zarringhalam, K.; Haspel, N., Patch-DCA: Improved Protein Interface Prediction by utilizing Structural Information and Clustering DCA scores. BioRxiv 2019, 656074.

29. Vreven, T.; Moal, I. H.; Vangone, A.; Pierce, B. G.; Kastritis, P. L.; Torchala, M.; Chaleil, R.; Jiménez-García, B.; Bates, P. A.; Fernandez-Recio, J., Updates to the integrated protein– protein interaction benchmarks: docking benchmark version 5 and affinity benchmark version 2. Journal of molecular biology 2015, 427, 3031–3041.

30. Berman, H. M.; Westbrook, J.; Feng, Z.; Gilliland, G.; Bhat, T. N.; Weissig, H.; Shindyalov, I. N.; Bourne, P. E., The protein data bank. Nucleic acids research 2000, 28, 235–242.

31. Lensink, M. F.; Wodak, S. J., Score_set: a CAPRI benchmark for scoring protein complexes. Proteins: Structure, Function, and Bioinformatics 2014, 82, 3163–3169.

32. Dolinsky, T. J.; Nielsen, J. E.; McCammon, J. A.; Baker, N. A., PDB2PQR: an automated pipeline for the setup of Poisson–Boltzmann electrostatics calculations. Nucleic acids research 2004, 32, W665–W667.

33. Wimley, W. C.; White, S. H., Experimentally determined hydrophobicity scale for proteins at membrane interfaces. Nature Structural and Molecular Biology 1996, 3, 842.

34. Goldenberg, O.; Erez, E.; Nimrod, G.; Ben-Tal, N., The ConSurf-DB: pre-calculated evolutionary conservation profiles of protein structures. Nucleic acids research 2008, 37, D323–D327.

35. Pupko, T.; Bell, R. E.; Mayrose, I.; Glaser, F.; Ben-Tal, N., Rate4Site: an algorithmic tool for the identification of functional regions in proteins by surface mapping of evolutionary determinants within their homologues. Bioinformatics 2002, 18, S71–S77.

36. Sanner, M. F.; Olson, A. J.; Spehner, J. C., Reduced surface: an efficient way to compute molecular surfaces. Biopolymers 1996, 38, 305–320.

37. Cock, P. J.; Antao, T.; Chang, J. T.; Chapman, B. A.; Cox, C. J.; Dalke, A.; Friedberg, I.; Hamelryck, T.; Kauff, F.; Wilczynski, B., Biopython: freely available Python tools for computational molecular biology and bioinformatics. Bioinformatics 2009, 25, 1422–1423.

38. Pedregosa, F.; Varoquaux, G.; Gramfort, A.; Michel, V.; Thirion, B.; Grisel, O.; Blondel, M.; Prettenhofer, P.; Weiss, R.; Dubourg, V., Scikit-learn: Machine learning in Python. Journal of machine learning research 2011, 12, 2825–2830.

39. Suhre, K.; Sanejouand, Y.-H., ElNemo: a normal mode web server for protein movement analysis and the generation of templates for molecular replacement. Nucleic acids research 2004, 32, W610–W614.

40. Suzuki, H.; Kawabata, T.; Nakamura, H., Omokage search: shape similarity search service for biomolecular structures in both the PDB and EMDB. Bioinformatics 2015, 32, 619–620.

41. Radermacher, M.; Ruiz, T., On cross-correlations, averages and noise in electron microscopy. Acta Crystallographica Section F: Structural Biology Communications 2019, 75, 12–18.

42. Meenan, N. A.; Sharma, A.; Fleishman, S. J.; MacDonald, C. J.; Morel, B.; Boetzel, R.; Moore, G. R.; Baker, D.; Kleanthous, C., The structural and energetic basis for high selectivity in a high-affinity protein-protein interaction. Proceedings of the National Academy of Sciences 2010, 107, 10080–10085.

43. Simon, A. J.; Zhou, Y.; Ramasubramani, V.; Glaser, J.; Pothukuchy, A.; Gollihar, J.; Gerberich, J. C.; Leggere, J. C.; Morrow, B. R.; Jung, C., Supercharging enables organized assembly of synthetic biomolecules. Nature chemistry 2019, 1.

44. Tan, P. H.; Sandmaier, B. M.; Stayton, P. S., Contributions of a highly conserved VH/VL hydrogen bonding interaction to scFv folding stability and refolding efficiency. Biophysical journal 1998, 75, 1473–1482.

45. Zhou, H.-X.; Qin, S., Interaction-site prediction for protein complexes: a critical assessment. Bioinformatics 2007, 23, 2203–2209.

46. de Vries, S. J.; Bonvin, A. M., How proteins get in touch: interface prediction in the study of biomolecular complexes. Current protein and peptide science 2008, 9, 394–406.

47. Panchenko, A. R.; Kondrashov, F.; Bryant, S., Prediction of functional sites by analysis of sequence and structure conservation. Protein science 2004, 13, 884–892.

48. Gong, X.; Wang, P.; Yang, F.; Chang, S.; Liu, B.; He, H.; Cao, L.; Xu, X.; Li, C.; Chen, W., Protein–protein docking with binding site patch prediction and network-based terms enhanced combinatorial scoring. Proteins: Structure, Function, and Bioinformatics 2010, 78, 3150–3155.

49. Xue, L. C.; Jordan, R. A.; Yasser, E. M.; Dobbs, D.; Honavar, V., DockRank: Ranking docked conformations using partner-specific sequence homology-based protein interface prediction. Proteins: Structure, Function, and Bioinformatics 2014, 82, 250–267.

50. De Vries, S. J.; Van Dijk, M.; Bonvin, A. M., The HADDOCK web server for data-driven biomolecular docking. Nature protocols 2010, 5, 883.

51. Cukuroglu, E.; Engin, H. B.; Gursoy, A.; Keskin, O., Hot spots in protein–protein interfaces: Towards drug discovery. Progress in biophysics and molecular biology 2014, 116, 165–173.

52. Jin, L.; Wang, W.; Fang, G., Targeting protein-protein interaction by small molecules. Annual review of pharmacology and toxicology 2014, 54, 435–456.

