## Supplementary Information for "Simplified geometric representations of protein structures identify complementary interaction interfaces"

1. diagnosticTestingSet.zip
   PDB, PQR, mutation rate, and confusion matrix files of the protein docking benchmark 5.0.

2. Normalmodepdbs.zip
   PDB files for normal mode analysis of PDBID: 1FQJ.

3. ProtenComplexBech5stats.xlsx
   Confusion matrix results for CAPRI and protein docking benchmark 5.0 sets.
