## Supplementary material for "Simplified geometric representations of protein structures identify complementary interaction interfaces": PDB, PQR, mutation rate, and confusion matrix files of the protein docking benchmark 5.0.: atmtypenumbers

# atmtypenumber 
# $Revision: 1.13 $
# Maps residue type and atom name pairs into Connolly ".atm" numeric codes
#  as used in MS and AMS, and into actual radius values
#
# Format: (blank lines and lines beginning with # are ignored)
#  Any number of blanks or tabs separate fields.
#
# Format of "radius" entries
# Field 1: keyword "radius"
# Field 2: atom number (key)
# Field 3: covalent bond distance taken from Connolly "ms.rad" file (unchecked)
# Field 4: atomic radius (angstroms) taken from Connolly "ms.explicit.rad" file
# Field 5: optional atomic radius for united-atom ("no hydrogens") approach
#  		taken from from Connolly "ms.rad" file
#
# By convention, radius entries come before atom type entries.
#
#
# Format of atom type entries:
# Field 1 : residue name pattern
# Field 2 : atom name pattern
# Field 3 : atom number (key value)
#
#
# Patterns are as in 'awk' or 'egrep', with the convention that '*' by itself
#  as a field means '.*', i.e., matches anything
#  and that underscores in the atom name represent blanks
# Note that ranges (such as '[1-4]' in 'P[1-4]') are by character, not numeric
#  so 'P[0-99]' does not do what you might expect.

# First match is used, so put catch-alls at the end.

# Adapted from file "protein.typ" by Michael Connolly, John Tainer,
# and Elizabeth Getzoff.
#  Michael Pique, Scripps Clinic
#
# $Log: atmtypenumbers,v $
# Revision 1.13  1997/09/09  03:33:05  mp
# Added some catch-alls at in for cofactors, updated RCS file
# from some intermediate edits done by Garrett but (apparently)
# never checked in. His version numbers up to 1.15 are subsumed in this 1.13.
#
# Revision 1.15  96/07/22  09:54:30  garrett
# Changed LYS/R to LYS/K - at the request of David S. Goodsell
# R is now present in only ARG,HIP/HIE/HID
#
# Revision 1.14  96/07/15  11:28:30  garrett
# Added the aromatic carbon atom type A in the "catch alls" at the end,
# and the phosphorus catch all, P.*
# for David S. Goodsell.
#
# Revision 1.13  96/07/08  13:03:29  garrett
# Added the non-standard atom types E (for charged oxygens) and
# R (for charged nitrogens) in ASP,GLU and ARG,LYS,HIP/HIE/HID respectively, 
# on behalf of David S. Goodsell.
#
# Revision 1.12  96/06/03  18:39:58  mp
# Allow digits in front of hydrogen & deut names
# 
# Revision 1.11  95/11/02  19:45:06  mp
# Added radius table (some values unchecked) and a few catch-alls
# 
# Revision 1.10  95/11/02  17:55:51  mp
# (13 May 1994): added missing VAL CB entry, added general-purpose C,O,N
# entries at end.
# 
# Revision 1.9  93/06/11  22:11:53  mp
# Added residue-type independent "CG", "CD", ... to deal with files
# where the sequence is not given.
# 
# Revision 1.8  93/05/19  23:01:03  mp
# Added "PO4" as synonym for PHO
# 
# Revision 1.7  93/05/19  18:11:17  mp
# Added deuterium completely same as hydrogen.
# 
# Revision 1.6  92/09/21  00:34:23  mp
# Added code for FE (not in heme), provisionally.
# 
# Revision 1.5  92/09/21  00:32:07  mp
# Added H2O for waters, allowed O.* in a water residue to match.
# Added "DOT" (23) and provisionally added MN MN (Manganese).
# 
# Revision 1.4  91/11/25  15:02:33  mp
# Corrected ASP OD1[AB] from previous update.
# 
# Revision 1.3  91/11/25  14:59:31  mp
# Added "A" and "B" as legal endings to ARG NH[12], GLU/ASP O[DE][12]
# 
# Revision 1.2  91/02/23  01:12:37  mp
# Corrections and additions to HIS family, especially HID-HIE interchange
# that had been incorrect since VAX version.  No longer insist that Cu and
# Zn be in residues of those names (allowing them to appear in HIS...)
# 
# Revision 1.1  90/02/27  12:34:46  mp
# Initial revision
# 
#
#   atom num dist  Rexplicit Runited-atom
radius    1  0.57    1.40
radius    2  0.66    1.40         1.60
radius    3  0.57    1.40
radius    4  0.70    1.54         1.70
radius    5  0.70    1.54         1.80
radius    6  0.70    1.54         2.00
radius    7  0.77    1.74         2.00
radius    8  0.77    1.74         2.00
radius    9  0.77    1.74         2.00
radius   10  0.67    1.74
radius   11  0.70    1.74         1.86
radius   12  1.04    1.80         1.85
radius   13  1.04    1.80  # P, S, and LonePairs
radius   14  0.70    1.54  # non-protonated nitrogens
radius   15  0.37    1.20  # H, D  hydrogen and deuterium
radius   16  0.70    0.00         1.50   # obsolete entry, purpose unknown
radius   17  3.50    5.00  # pseudoatom - big ball
radius   18  1.74    1.97  # Ca calcium
radius   19  1.25    1.40  # Zn zinc    (traditional radius)
radius   20  1.17    1.40  # Cu copper  (traditional radius)
radius   21  1.45    1.30  # Fe heme iron
radius   22  1.41    1.49  # Cd cadmium
radius   23  0.01    0.01  # pseudoatom - tiny dot
radius   24  0.37    1.20         0.00   # hydrogen vanishing if united-atoms
radius   25  1.16    1.24  # Fe not in heme
radius   26  1.36    1.60  # Mg magnesium
radius   27  1.17    1.24  # Mn manganese
radius   28  1.16    1.25  # Co cobalt
radius   29  1.17    2.15  # Se selenium
radius   30  3.00    3.00  # obsolete entry, original purpose unknown
radius   31  1.15    1.15  # Yb ytterbium +3 ion --- wild guess only
radius   38  0.95    1.80  # obsolete entry, original purpose unknown
#
# note that the metal values are UNCHARGED radii, see
#  http://www.shef.ac.uk/chemistry/web-elements for info  - Mike Pique

# Hydrogens - not differentiated here
*    [0-9]*H.*    15
*    [0-9]*D.*    15

# Waters and confusingly-named metals
WAT|HOH|H2O|DOD|DIS  O.*        2
CA   CA      18 
CD   CD      22
*    CD__    22

# Atoms that are sometimes named like mainchain atoms but aren't really
#  get caught here
ACE  CA	      9

# Mainchain atoms - invariant by residue/nucleotide type
*    N        4
*    CA       7
*    C       10
*    O        1
*    P       13

# CB - C beta
ALA  CB       9
ILE|THR|VAL  CB   7
*    CB       8

# CG - C gamma
ASN|ASP|ASX|HIS|HIP|HIE|HID|HISN|HISL|LEU|PHE|TRP|TYR  CG      10
ARG|GLU|GLN|GLX|MET  CG       8
LEU  CG       7
*    CG       8


# Other amino acid residues listed by residue type
# note the "question mark" matches zero or one occurances of pattern
GLN  O1       3
GLN  O2       3
ACE  CH3      9
ARG  CD       8
ARG  NE       4
ARG  RE       4
ARG  CZ      10
ARG  NH[12][AB]?  5
ARG  RH[12][AB]?  5
ASN  OD1      1
ASN  ND2      5
ASN  AD1      3
ASN  AD2      3
ASP  OD[12][AB]? 3
ASP  ED[12][AB]? 3
ASX  OD1[AB]? 1
ASX  ND2      5
ASX  AD1      3
ASX  AD2      3
ASX  OD2      3
CYS|MET  LP[12]     13
CY[SXM]  SG      13
CYH  SG      12
GLU  OE[12][AB]?    3
GLU  EE[12][AB]?    3
GLU|GLN|GLX  CD      10
GLN  OE1      1
GLN  NE2      5
GLN|GLX  AE[12]      3

# His and relatives
# There are 4 kinds of HIS rings: HIS (no protons), HID (proton on Delta),
#   HIE (proton on epsilon), and HIP (protons on both)
# Protonated nitrogens are numbered 4, else 14
# HIS is treated here as the same as HIE
# 
# HISL is a deprotonated HIS (the L means liganded)

HIS|HID|HIE|HIP|HISL	CE1|CD2		11
HIS|HIE|HISL 		ND1		14
HID|HIP			ND1		 4
HID|HIP			RD1		 4
HIS|HIE|HIP		NE2		 4
HIS|HIE|HIP		RE2		 4
HID|HISL		NE2		14
HID|HISL		RE2		14
HIS|HID|HIP|HISD	A[DE][12]	 4

ILE  CG1      8
ILE  CG2      9
ILE  CD|CD1      9

LEU  CD1      9
LEU  CD2      9
LYS  C[GDE]       8
LYS  NZ       6
LYS  KZ       6
MET  SD      13
MET  CE       9

PHE  C[DE][12]     11
PHE  CZ      11

PRO|CPR  C[GD]       8
CSO  SE       9
CSO  SEG      9
CSO  OD1      3
CSO  OD2      3
SER  OG       2
THR  OG1      2
THR  CG2      9
TRP  CD1     11
TRP  CD2     10
TRP  CE2     10
TRP  NE1      4
TRP  CE3     11
TRP  CZ2     11
TRP  CZ3     11
TRP  CH2     11
TYR  C[DE][12]     11
TYR  CZ      10
TYR  OH       2
VAL  CG1      9
VAL  CG2      9

# catch common atom names for non-standard residue names
*    CD       8
*    CE       8

#
# check these next two with JAT & EDG mp
# (numbering up to 7 is on suggestion of Dave Stout 20 Feb 90 mp)
FS[34]  FE[1-7]     21
FS[34]  S[1-7]      13
FS3  OXO      1
FEO  FE1     21
FEO  FE2     21


HEM  O1       1
HEM  O2       1
HEM  FE      21
HEM  CH[A-D] 11
HEM  N[A-D]  14
HEM  N_[A-D] 14
HEM  C[1-4][A-D]     10
HEM  CM[A-D]  9
HEM  C[AB][AD]     8
HEM  CG[AD]  10
HEM  O[12][AD]      3
HEM  C[AB][BC]     11
HEM  OH2      2
AZI  N[123]  14
MPD  C1       9
MPD  C2      10
MPD  C3       8
MPD  C4       7
MPD  C5       9
MPD  C6       9
MPD  O7       2
MPD  O8       2
SO4|SUL  S   13
SO4|SUL  O[1234]  3
PO4|PHO  O[1234]  3
PC   O4       3    
PC   P1      13    
PC   O[123]       3    
PC   C[12]    8    
PC   N1      14    
PC   C[345]   9    

# Special giant atom (big ball)
BIG  BAL     17

# Special tiny atom (point)
POI  POI     23
DOT  DOT     23

# Metals:

# Heroes of SOD
*   CU      20
*   ZN      19
# Note 24 and 25 have not been OKed by John & Libby - MP, September 1992
*   MN      24
# This free FE is not the same number as heme FE
*   FE      25
# These are metals, considered uncharged so use at your own risk
*   MG      26
*   MN      27
*   CO      28
*   SE      29
*   YB      31

# Unknown soldiers:

# FMN is cofactor Flavin mononucleotide
FMN  N1       4
FMN  C[2478] 10
FMN  O2       1
FMN  N3      14
FMN  O4       1
FMN  C[459]A 10
FMN  N5       4
FMN  C[69]   11
FMN  C[78]M   9
FMN  N10      4
FMN  C10     10
FMN  C[12345]\* 8
FMN  O[234]\* 2
FMN  O5\*     3
FMN  OP[1-3]  3

ALK|MYR  OT1  3
ALK|MYR  C01 10
ALK  C16      9
MYR  C14      9
ALK|MYR  C.*  8

# Catch-alls

*    SEG      9
*    OXT      3
*    OT.*      3
*    E.*       3
*    S.*      13
*    C.*	7
# A is for aromatic carbons - GMM/DSG/AJO.
*    A.*       11
*    O.*	1
*    N.*	4
*    R.*	4
*    K.*	6
# PB might be Pb (lead) but its academic... Many PDB files have named P atoms
*    P[A-D]   13
# Added even more general phosphorus - GMM/DSG/AJO.
*    P.*       13

# further catch-alls for HETATM entries, e.g. AC1*, AO5, PC14, PO10 (MP 199709)
FAD|NAD|AMX|APU	.O.*	1
FAD|NAD|AMX|APU	.N.*	4
FAD|NAD|AMX|APU	.C.*	7
FAD|NAD|AMX|APU	.P.*	13
FAD|NAD|AMX|APU	.H.*	15
