## Supplementary material for "Simplified geometric representations of protein structures identify complementary interaction interfaces": PDB, PQR, mutation rate, and confusion matrix files of the protein docking benchmark 5.0.: diagnosticTesting.py

```
import argparse
import numpy as np
import operator
import MorphProt
import pandas as pd
from MorphProt import proteinShell
from MorphProt import physicalProp
from MorphProt import shapeProcessing
from MorphProt import cubeFaceData
from MorphProt import boxDictionary
from MorphProt import faceBuilder
from MorphProt import toSphere
from MorphProt import topCubeScore
from MorphProt import topSphereScore
from MorphProt import mapBack
import csv
import Bio
from scipy.spatial import distance
from heapq import nsmallest
from heapq import nlargest

def pdbpqr_parse(name, file):
    pdb_parser = Bio.PDB.PDBParser(QUIET = True)
    ref_structure = pdb_parser.get_structure(name, file)
    model = ref_structure[0]
    print("PDB/PQR file is being parsed...")
    return model

def protein_coord_extract(name, file):
    pdb_parser = Bio.PDB.PDBParser(QUIET = True)
    ref_structure = pdb_parser.get_structure(name, file)
    model = ref_structure[0]
    atom_data = []
    for chain in model:
        for residue in chain:
            y = Bio.PDB.is_aa(residue)
            if y == True:
                aa_res = residue
                res_num = aa_res.get_id()[1]
                for atom in aa_res:
                    XYZ = atom.get_vector()
                    X = XYZ[0]
                    Y = XYZ[1]
                    Z = XYZ[2]
                    value = [res_num, X, Y, Z]
                    atom_data.append(value)
    return atom_data

def table_content(name):
    # open the file and read the contents within and keep the formatting of the table
    filename = '/home/cns-mccafferty/coyote/Electrostatics/benchmark5/validation/interface_distances/PR_files/diagnosticTestingSet/' + name 
    #filename = '/stor/home/ghoogerb/anaconda3/Marcotte/caitie/MorphProt-master/working/consFiles/' + name + '.txt'
    content = pd.read_table(filename, skiprows=2, usecols=[0,1,2,3,5,6,9,12,13], names = ["POS", "SEQ", "3LATOM", "SCORE", "COLOR", "CONFIDENCE INTERVAL", "CONFIDENCE INTERVAL COLORS", "MSA DATA", "RESIDUE VARIETY"])
    return content
def renumbering(name):
    table = table_content(name)
    df = pd.DataFrame(table)
    for row in range(len(table)):
        atom_seq = table.get_value(row, "3LATOM")
        atom = ''
        atom_num = atom.join(x for x in atom_seq if x.isdigit())
        position = table.get_value(row, "POS")
        # compare the 3LATOM atom position with the POS atom position
        try:
            if int(atom_num) == position:
                continue
            elif int(atom_num) != position:
                df.loc[[row], ['POS']] = int(atom_num)
        except ValueError:
            continue
    return df
def renumbering_dashes(name):
    # renumbering the rows with '-' as 3LATOM
    dataframe = renumbering(name)
    for row in range(len(dataframe)):
        if row >= 1 and (dataframe.get_value(row, "POS") - dataframe.get_value(row-1, "POS")) > 1:
            while (row > 0):
                # loop to change all the lines above the row found to have a difference greater than 1
                dataframe.loc[[row-1], ['POS']] = int(dataframe.get_value(row, "POS") - 1)
                row -= 1
        else:
            continue
    for rows in range(len(dataframe)):
        if rows >= 1 and (dataframe.get_value(rows-1, "POS") - dataframe.get_value(rows, "POS")) > 1:
            while (rows < len(dataframe)):
                # loop to change all the lines after the row found to have a difference greater than 1
                dataframe.loc[[rows], ['POS']] = int(dataframe.get_value(rows-1, "POS") + 1)
                rows += 1
    return dataframe
def cleaning_cons(name):
    # function to remove '*' from column COLOR
    cons_data = renumbering_dashes(name)
    for row in range(len(cons_data)):
        color = cons_data.get_value(row, "COLOR")
        if "*" in color:
            stripped_color = color.replace("*", "")
            cons_data.loc[[row], ['COLOR']] = stripped_color
        else:
            continue
    return cons_data
def saving_cons(name):
    try:
        cleaned_cons = cleaning_cons(name)
    except TypeError:
        cleaned_cons = renumbering_dashes(name)
    #cons = cleaned_cons.to_csv(r'/stor/home/ghoogerb/anaconda3/Marcotte/caitie/MorphProt-master/working/consFiles/consFiles_modified/' + name + '_cons.txt', sep='\t', index= False, mode='a')
    return cleaned_cons

def protein_cube_transform(name, pdb, pqr, attribute, mut_rate=None):

    # gives list of surface residues
    model = MorphProt.proteinShell.pdbpqr_parse(name, pqr)
    protein_surface = MorphProt.proteinShell.protein_shell(model)
    
    if attribute == str(2):
        # gets mutation rates for residues
        # mutation_rate_pd = pd.read_table(mut_rate, skiprows=1, usecols=[0,1,2,3,4,5,6,9,12], names = ["POS", "SEQ", "3LATOM", "SCORE", "COLOR", "CONFIDENCE INTERVAL", "CONFIDENCE INTERVAL COLORS", "MSA DATA", "RESIDUE VARIETY"])
        mutation_rate_pd = saving_cons(mut_rate)
        mut_rate_dict = dict(zip(mutation_rate_pd.POS, mutation_rate_pd.COLOR))
        print(mut_rate_dict)
        if len(str(mut_rate_dict[mutation_rate_pd.POS[0]])) >1:
            mut_rate_dict = {key: item.strip()[0] for key, item in mut_rate_dict.items()}
        else:
            mut_rate_dict = dict((k,str(v)) for k,v in mut_rate_dict.items())
        
        # Inserts properties into the, the results gives max/min coords, all coords, and residue information w hydropathy
        cuboid = MorphProt.physicalProp.cuboid_dimensions(model, protein_surface, mut_rate_dict)

        # the begining of the transformation where a k-menas is performed to divide the protein into 6 faces, returns the projected 2D faces
        xyz_list = MorphProt.shapeProcessing.protein_processing(pdb, cuboid)

        # gives the 2D projection coordinates paired with their corresponding properties
        coord_prop_list = MorphProt.cubeFaceData.coord_charges(xyz_list, cuboid)
            
    else:
        
        # Inserts properties into the, the results gives max/min coords, all coords, and residue information w hydropathy
        cuboid = MorphProt.physicalProp.cuboid_dimensions(model, protein_surface)

        # the begining of the transformation where a k-menas is performed to divide the protein into 6 faces, returns the projected 2D faces
        xyz_list = MorphProt.shapeProcessing.protein_processing(pdb, cuboid)

        # gives the 2D projection coordinates paired with their corresponding properties
        coord_prop_list = MorphProt.cubeFaceData.coord_charges(xyz_list, cuboid)

    return xyz_list, cuboid, coord_prop_list

def cube_face_assembly(xyz_list, coord_prop_list, resolution, shuffle, attribute, shape):    
    
    # divides up the coordinates into their faces
    prot_face_list = []
    for i in range(6):
        prot_face = MorphProt.boxDictionary.box_dictionary(xyz_list[i], coord_prop_list[i], resolution, shuffle)
        prot_face_list.append(prot_face)

    # produces 6 matrices with average property values, given the property being considered
    prot = MorphProt.faceBuilder.face_normalization(prot_face_list, attribute, shape)
    
    return prot_face_list, prot

def cross_corr_analysis(prot1_face_assembly, prot2_face_assembly, prot1, prot2, attribute, shape):
    if shape == 'cube':
        score_matrix = MorphProt.topCubeScore.cube_corr(prot1_face_assembly[1], prot2_face_assembly[1], attribute)
        top_scores = MorphProt.topCubeScore.top_cube_corr(score_matrix, attribute)
                
        if attribute == '0' or '2':
            score_key = max(top_scores, key=top_scores.get)
            score_loc = top_scores[score_key][1]
            
        if attribute == '1':
            score_key = min(top_scores, key=top_scores.get)
            score_loc = top_scores[score_key][1]
    return score_key, top_scores[score_key][0], map_to_protein1, map_to_protein2

def residue_id_extract(atom_coords_list):    
    atom_id = []
    for i in atom_coords_list:
        atom_id.append(i[1])

    atom_id_unique = sorted(set([i for i in atom_id if atom_id.count(i)>2]))
    return atom_id_unique

def model_data_extract(modelTrue):
    atom_data = []
    for chain in modelTrue:
        for residue in chain:
            y = Bio.PDB.is_aa(residue)
            if y == True:
                aa_res = residue
                res_num = aa_res.get_id()[1]
                for atom in aa_res:
                    XYZ = atom.get_vector()
                    X = XYZ[0]
                    Y = XYZ[1]
                    Z = XYZ[2]
                    value = [res_num, X, Y, Z]
                    atom_data.append(value)
    return atom_data

def true_interface(model1_atom_data, model2_atom_data):
    atomic_distances = {}
    model1_true_interface = []
    model2_true_interface = []
    for i in range(len(model1_atom_data)):
        for k in range(len(model2_atom_data)):
            key = str(model2_atom_data[k][0]) + '_' + str(model1_atom_data[i][0]) + '_' + str(i)
            value = distance.euclidean(model2_atom_data[k][1:], model1_atom_data[i][1:])
            if value <= 10:
                model1_true_interface.append(model1_atom_data[i][0])
                model2_true_interface.append(model2_atom_data[k][0])
    return model1_true_interface, model2_true_interface

def Main(name1, pdb1, pqr1, mutrate1, name2, pdb2, pqr2, mutrate2, attribute, shape):
    prot1 = protein_cube_transform(name1, pdb1, pqr1, attribute, mut_rate=mutrate1)
    prot2 = protein_cube_transform(name2, pdb2, pqr2, attribute, mut_rate=mutrate2)
    prot1_face_assembly = cube_face_assembly(prot1[0], prot1[2], resolution, 'off', attribute, shape)
    prot2_face_assembly = cube_face_assembly(prot2[0], prot2[2], resolution, 'off', attribute, shape)

    if attribute == '0':
        outfile = '_hydropathy'
    if attribute == '1':
        outfile = '_charge'
    if attribute == '2':
        outfile = '_mutrate'

    score_matrix = MorphProt.topCubeScore.cube_corr(prot1_face_assembly[1], prot2_face_assembly[1], attribute)
    top_scores = MorphProt.topCubeScore.top_cube_corr(score_matrix, attribute)

    # extraction of true interface
    model1True = pdbpqr_parse(name1, pdb1True)
    model2True = pdbpqr_parse(name2, pdb2True)

    model1_atom_data = model_data_extract(model1True)
    model2_atom_data = model_data_extract(model2True)

    model1_true_interface, model2_true_interface = true_interface(model1_atom_data, model2_atom_data)
    model1_true_interface = list(set(model1_true_interface))
    model2_true_interface = list(set(model2_true_interface))
    
    # calculates the smallest scores for charge prediction
    if attribute == '1':
        scores ={} # max scores for charge
        for k in score_matrix:
            bottom_three = nsmallest(3,[item for sublist in score_matrix[k].tolist() for item in sublist])
            for a,b in enumerate(bottom_three):
                scores[k + (a,)] = [bottom_three[a], np.unravel_index(np.argmin(score_matrix[k]), score_matrix[k].shape)]
        lowest = dict(sorted(scores.items(), key=operator.itemgetter(1), reverse=False)[:10])
        score_keys = list(lowest.keys())
        score_locs = [scores[i][1] for i in score_keys]
    
    # extracts maximum scores for hydropathy and mutation rate
    if attribute == '0' or '2':
        scores ={} # max scores for hydropathy
        for k in score_matrix:
            top_three = nlargest(3,[item for sublist in score_matrix[k].tolist() for item in sublist])
            for a,b in enumerate(top_three):
                scores[k + (a,)] = [top_three[a], np.unravel_index(np.argmax(score_matrix[k]), score_matrix[k].shape)]
        largest = dict(sorted(scores.items(), key=operator.itemgetter(1), reverse=True)[:10])
        score_keys = list(largest.keys())
        score_locs = [scores[i][1] for i in score_keys]

    # calculates true positive and false positive per prediction
    truePositives = []
    falsePositives = []
    for i in score_keys:
        out_matrix = MorphProt.mapBack.matrix_mapping(prot1_face_assembly[1][i[0]], prot2_face_assembly[1][i[1]], scores[i][1])
        matching_atomic_coords1 = MorphProt.mapBack.matching_atomic_coords(out_matrix[0], prot1_face_assembly[0][i[0]], attribute)
        matching_atomic_coords2 = MorphProt.mapBack.matching_atomic_coords(out_matrix[1], prot2_face_assembly[0][i[1]], attribute)
        map_to_protein1 = MorphProt.mapBack.cube_map_to_protein(prot1[1], matching_atomic_coords1, i[0])
        map_to_protein2 = MorphProt.mapBack.cube_map_to_protein(prot2[1], matching_atomic_coords2, i[1])
        protein1_id_extract = residue_id_extract(map_to_protein1)
        protein2_id_extract = residue_id_extract(map_to_protein2)
        truePositives.append(len(set(model1_true_interface) & set(protein1_id_extract)) + len(set(model2_true_interface) & set(protein2_id_extract)))
        falsePositives.append(len(protein1_id_extract) + len(protein2_id_extract))
    
    # calculates diagnostics from true interface and predicted interface for the top 10 predictions
    TP = sum(truePositives)
    interface = (len(model1_true_interface) + len(model2_true_interface))*10
    FN = interface - TP
    FP = sum(falsePositives) - TP
    precision = TP/(TP+FP)
    recall = TP/(TP+FN)
    F1 = (2 * precision * recall)/(precision + recall)
    
    with open(name1[:-2] + outfile + '_diagnostic.txt', 'a') as f:
        f.write(str(TP) + '\n')
        f.write(str(precision)+ '\n')
        f.write(str(recall)+ '\n')
        f.write(str(F1)+ '\n')

parser = argparse.ArgumentParser(description='Transform PDB structure to cubes to evaluate protein-protein intercations.')
parser.add_argument('name1', help='input protein l name for the structure of interest.')
parser.add_argument('name2', help='input protein 2 name for the structure of interest.')
parser.add_argument('attribute', metavar='att', help='physical property of protein that is beng studied, 0 for hydropathy, 1 for charge, and 2 for mutation rate.')
parser.add_argument('--shuffle', metavar='-s',  nargs='?', default='off', help='shuffles the boxes to test for different score minimums')
parser.add_argument('--resolution', metavar='-r',  nargs='?', default=5, help='the size, in angstroms, of the bins in each protein face')

args = parser.parse_args()

shape = 'cube'
resolution = args.resolution
pdb1True = args.name1 + '_u.pdb'
pdb2True = args.name2 + '_u.pdb'
pdb1 = args.name1 + '_u.pdb'
pqr1 = args.name1 + '_u.pqr'
mutrate1 = args.name1 + '_cons.txt'
pdb2 = args.name2 + '_u.pdb'
pqr2 = args.name2 + '_u.pqr'
mutrate2 = args.name2 + '_cons.txt'
attribute = args.attribute

Main(args.name1, pdb1, pqr1, mutrate1, args.name2, pdb2, pqr2, mutrate2, attribute, shape)
```
