## Supplementary figures and images for "Simplified geometric representations of protein structures identify complementary interaction interfaces"

### 1ATN_l_u.pdf

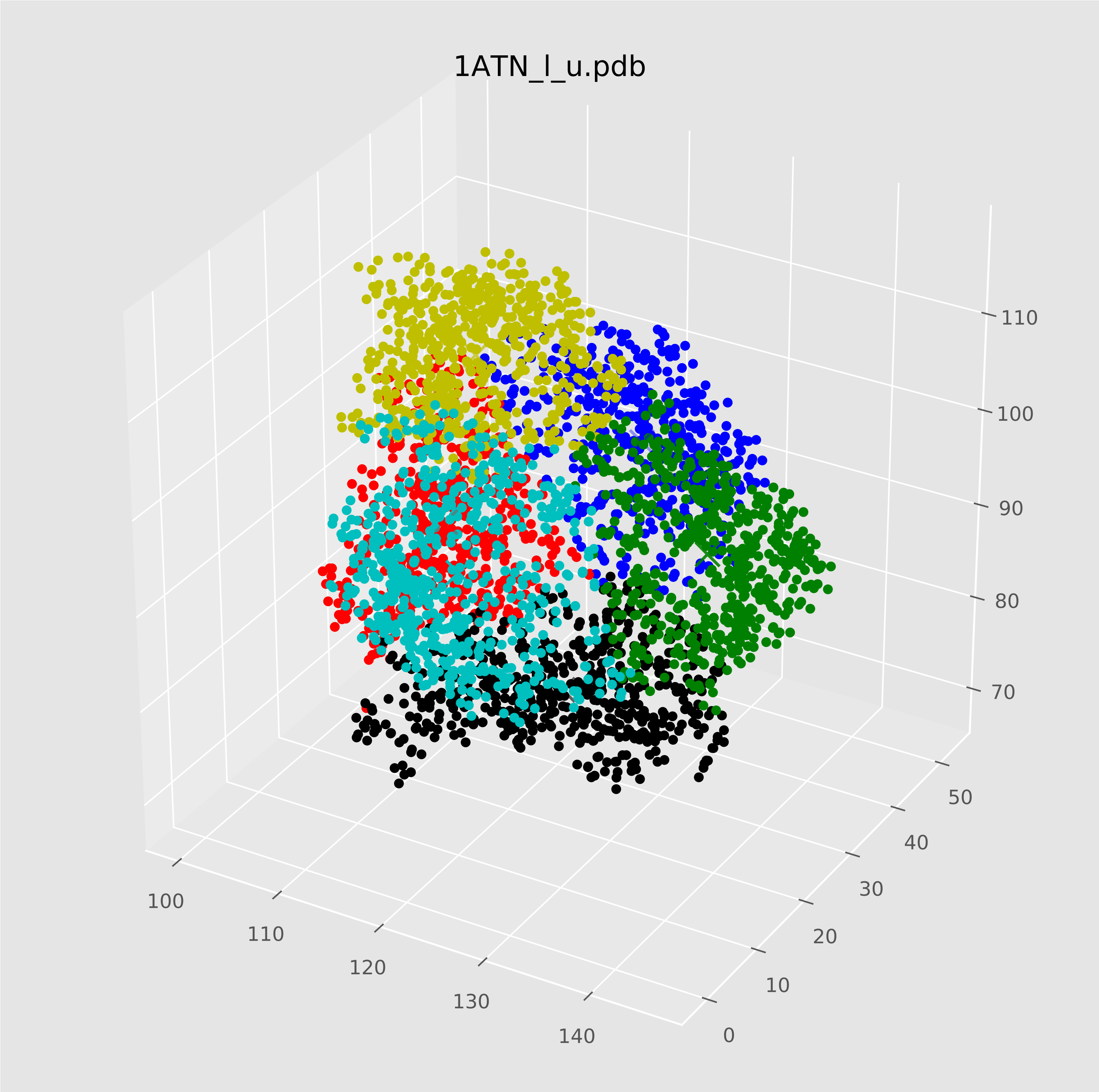

### 1ATN_r_u.pdf

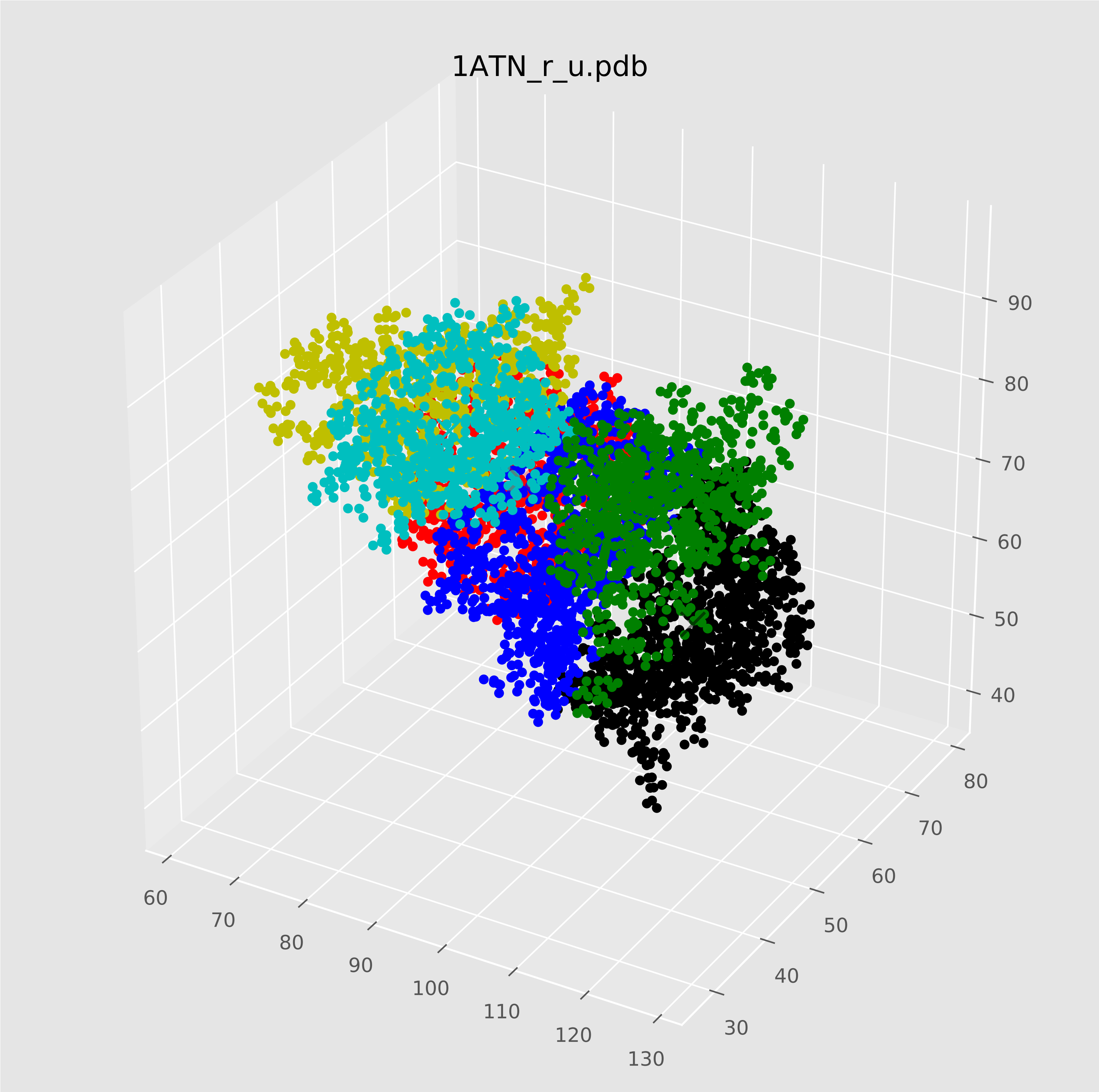

### 1D6R_l_u.pdf

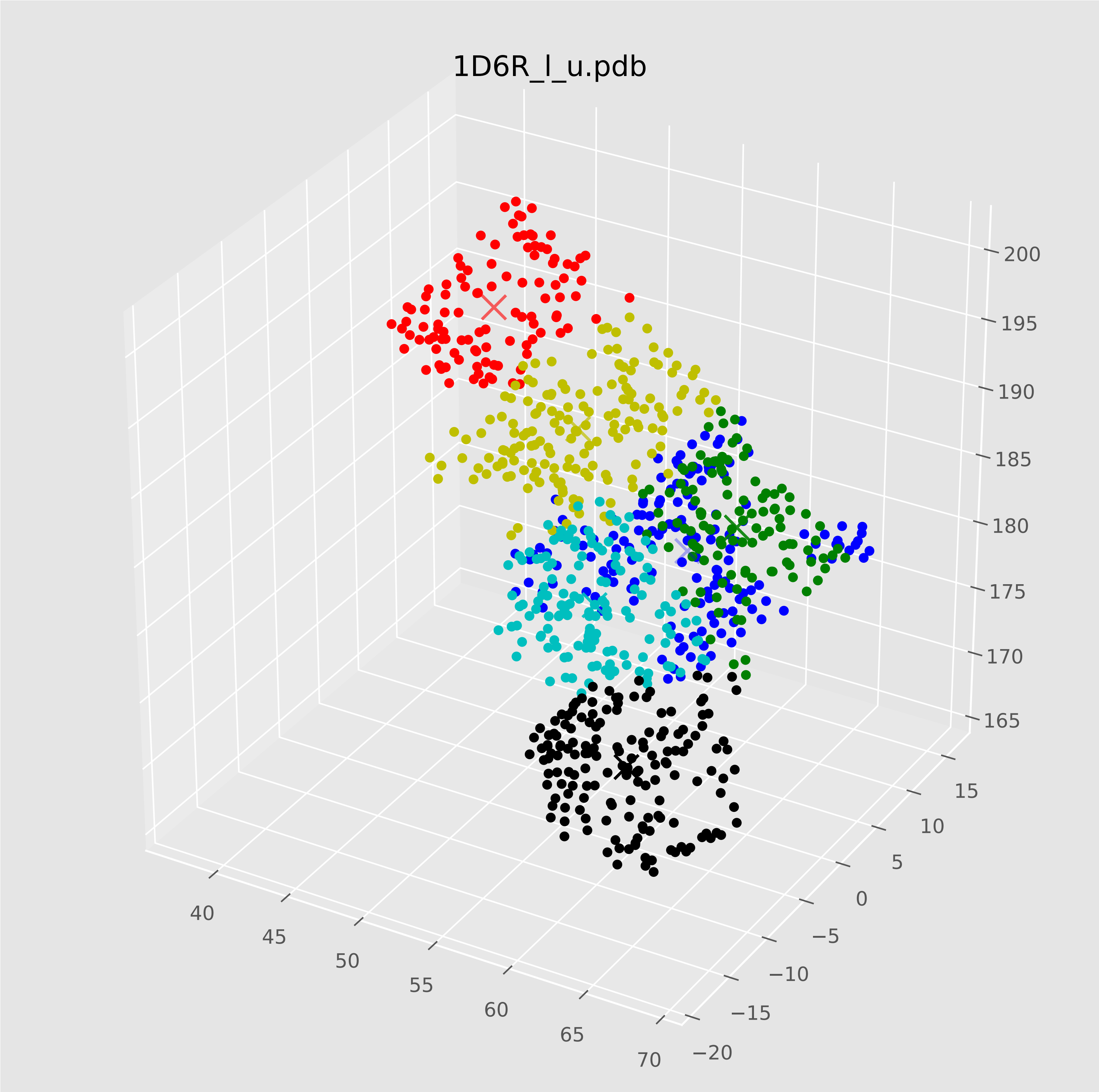

### 1D6R_r_u.pdf

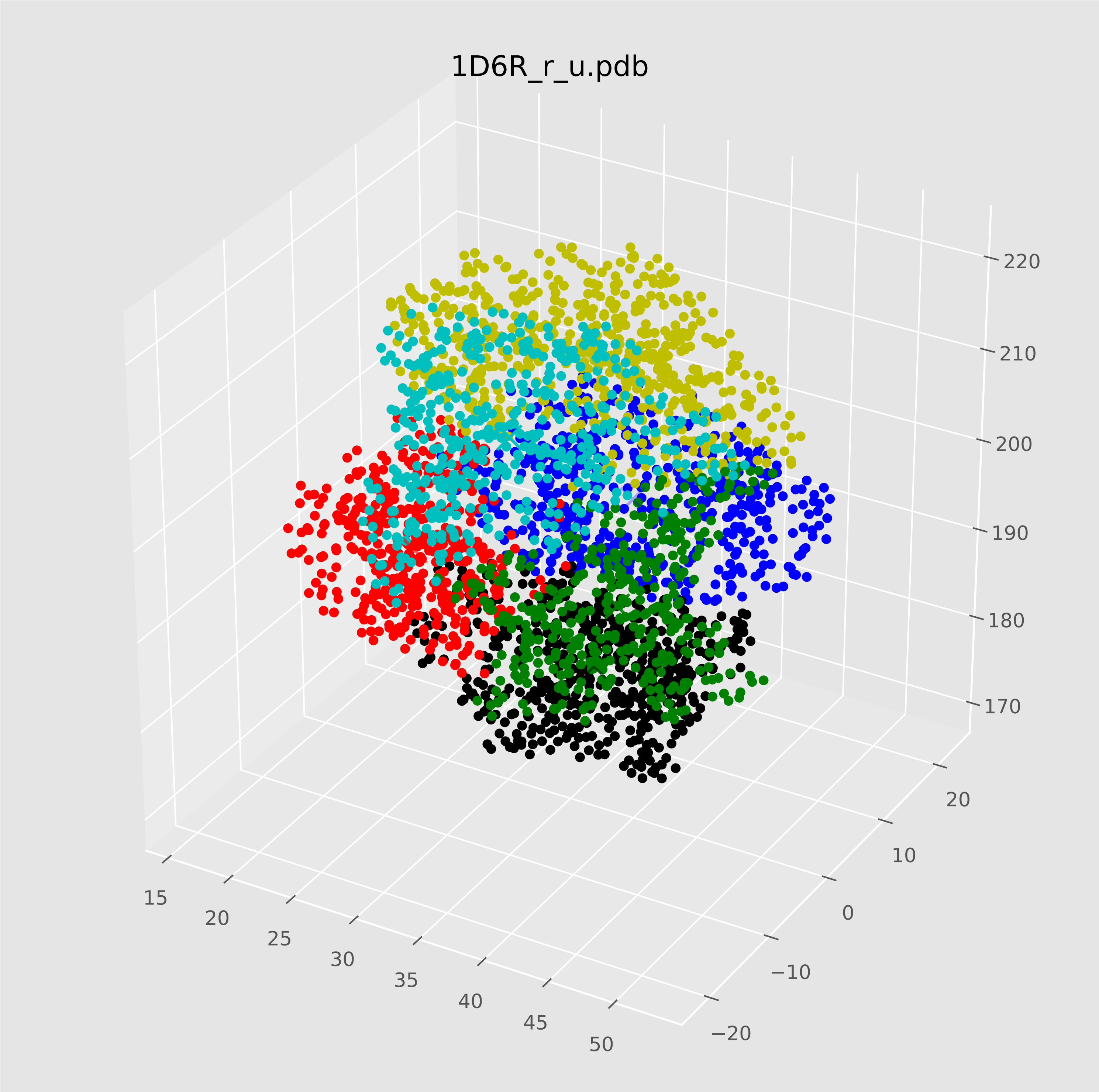

### 1GRN_l_u.pdf

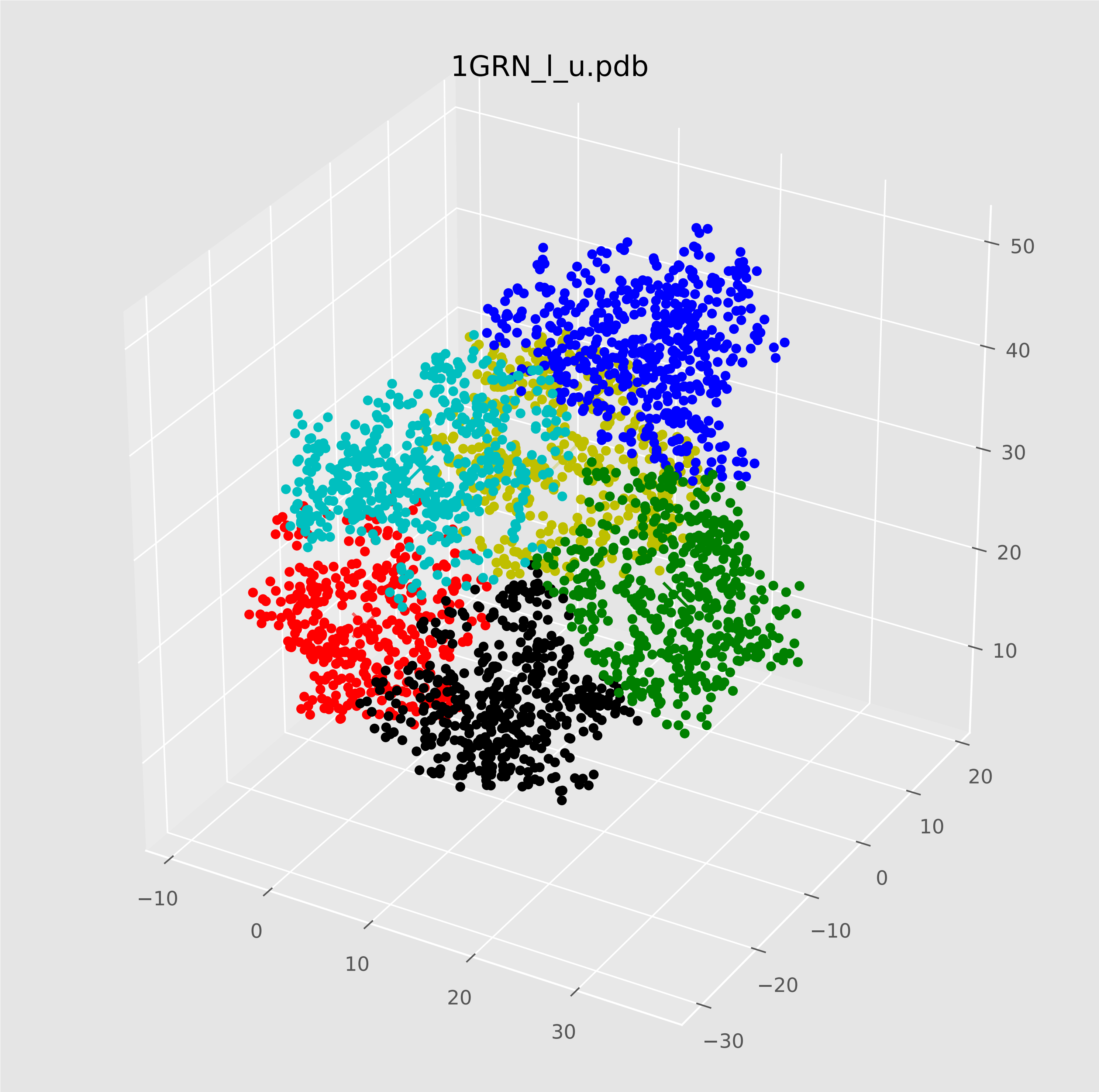

### 1GRN_r_u.pdf

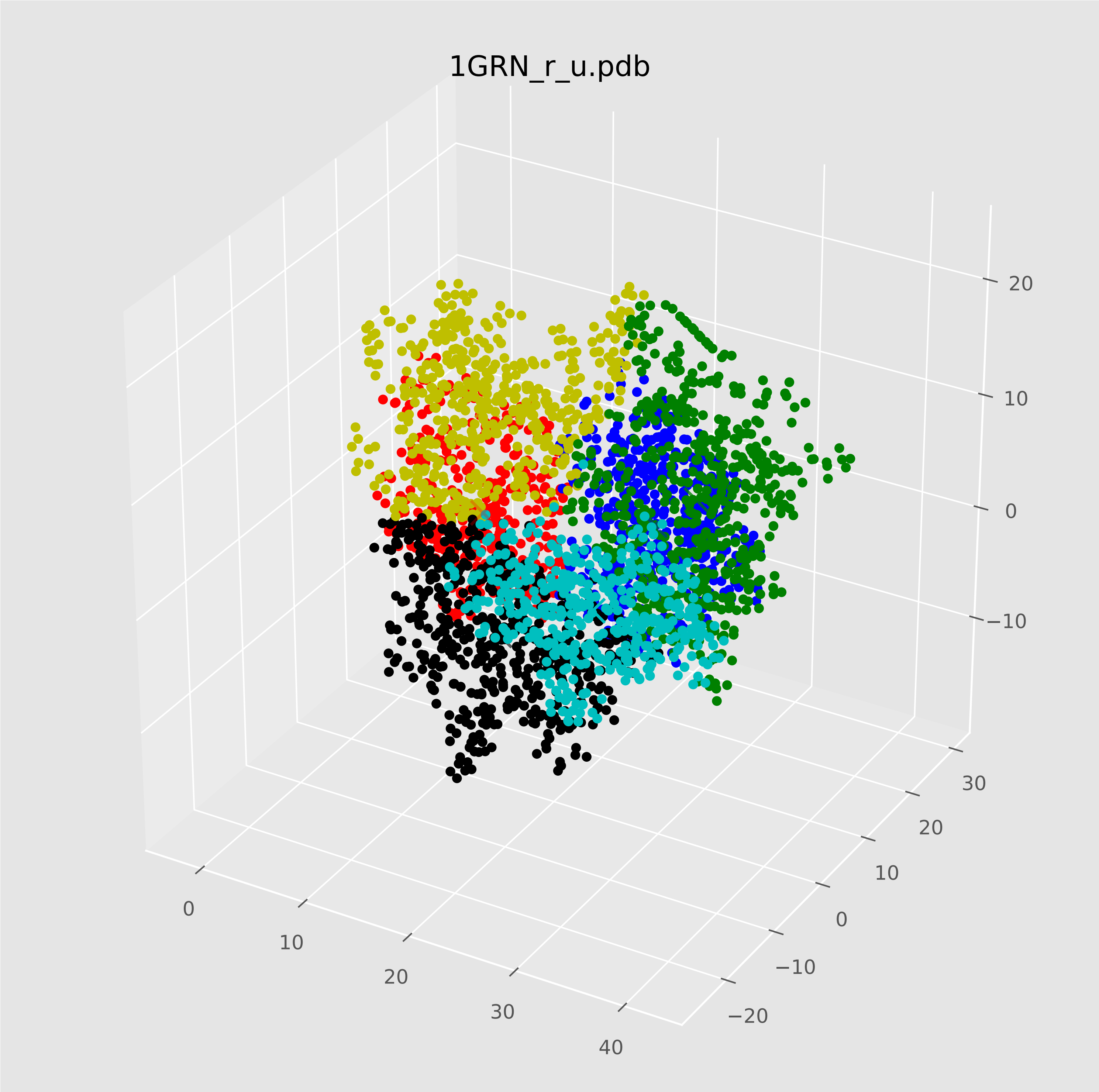

### 1KTZ_l_u.pdf

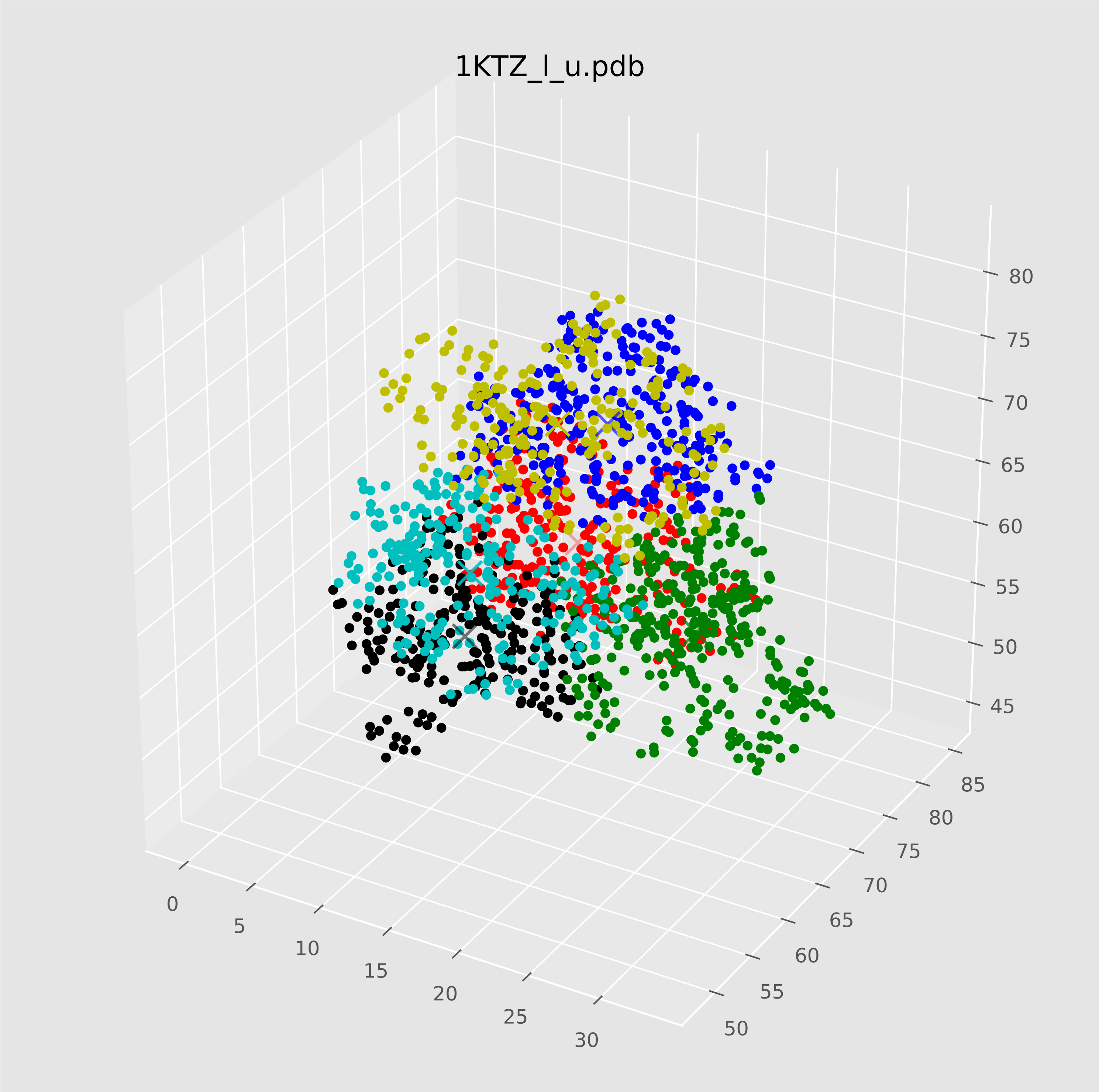

### 1KTZ_r_u.pdf

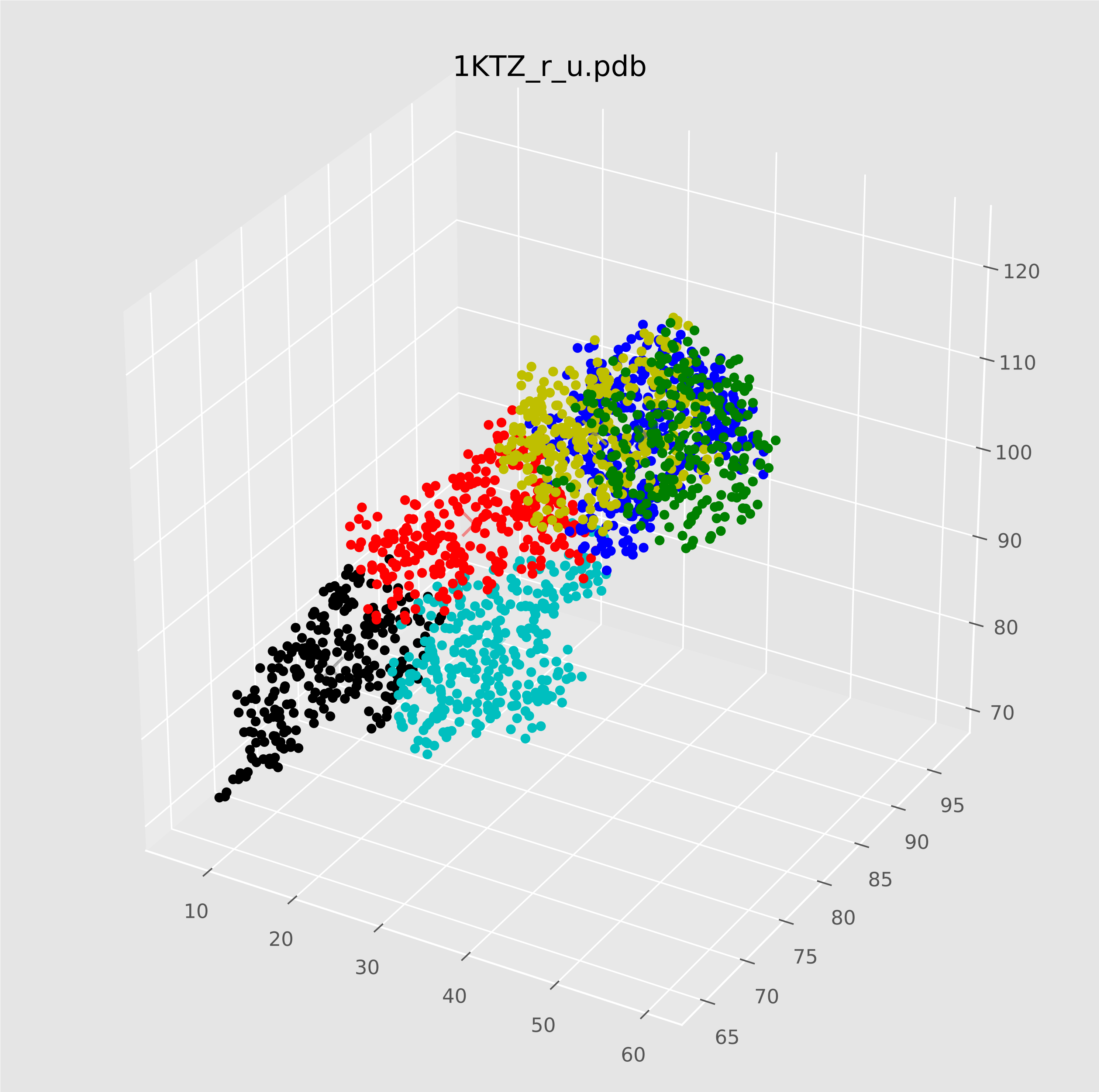

### 1KXP_l_u.pdf

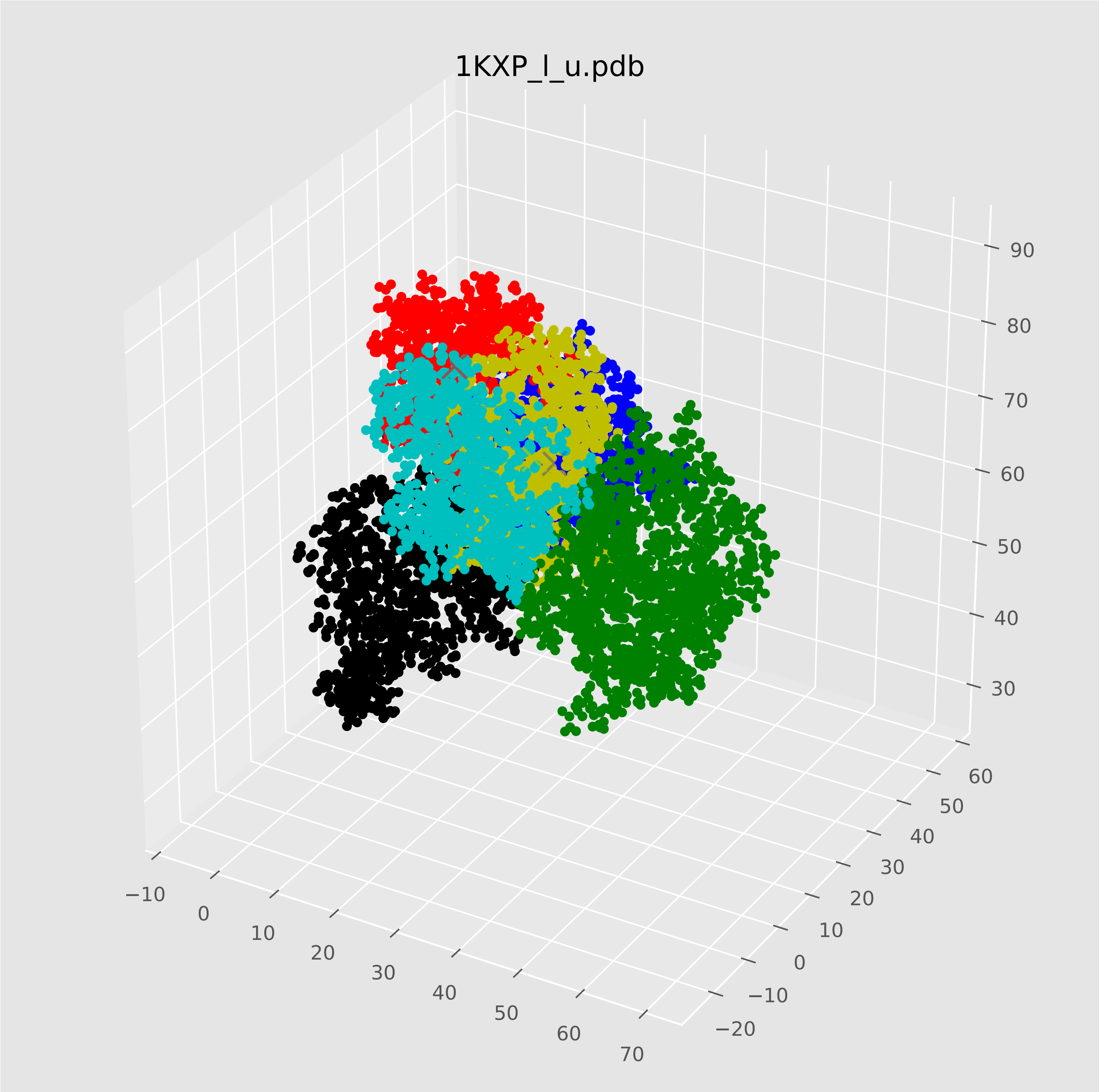

### 1KXP_r_u.pdf

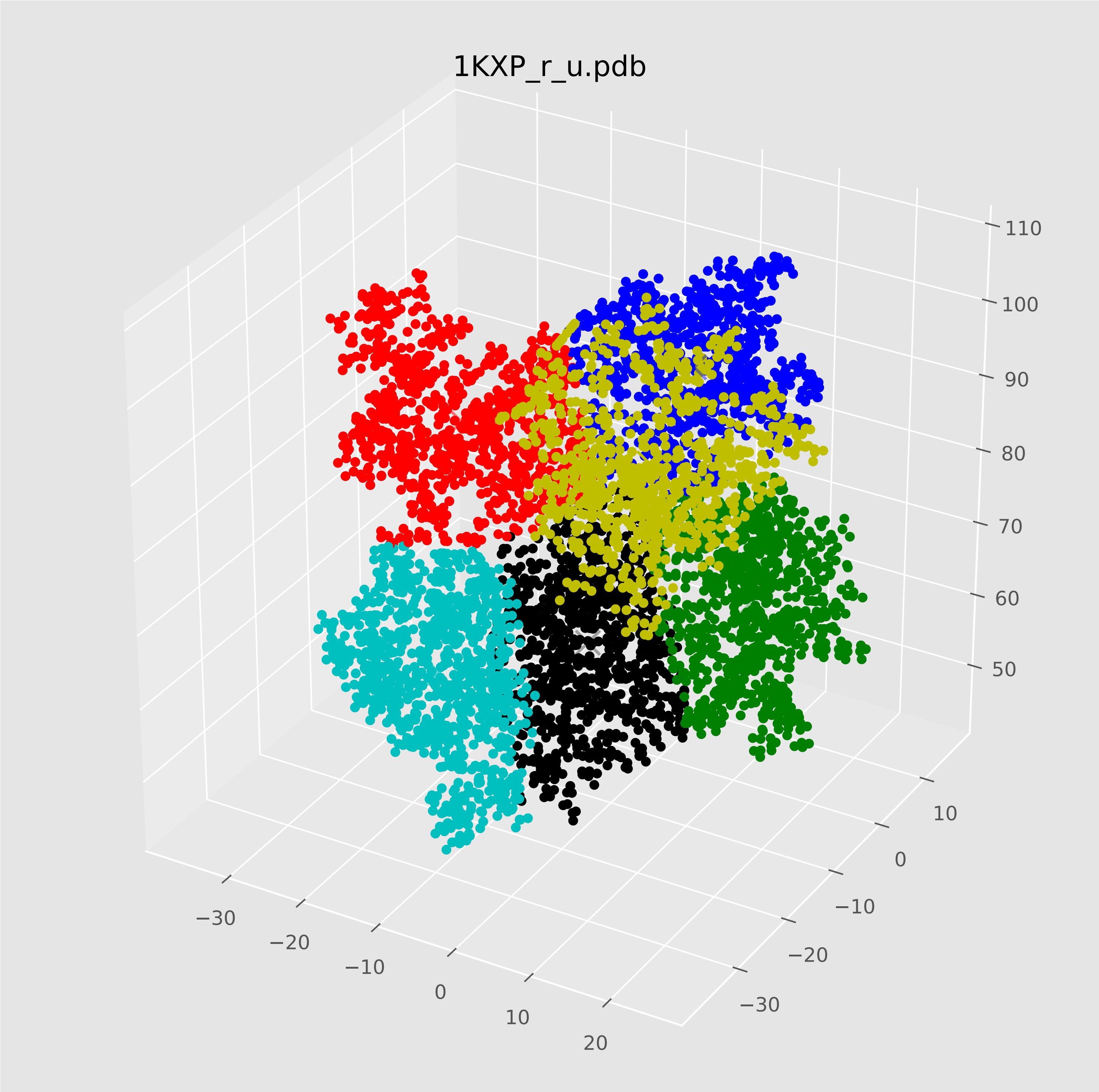

### 1KXQ_l_u.pdf

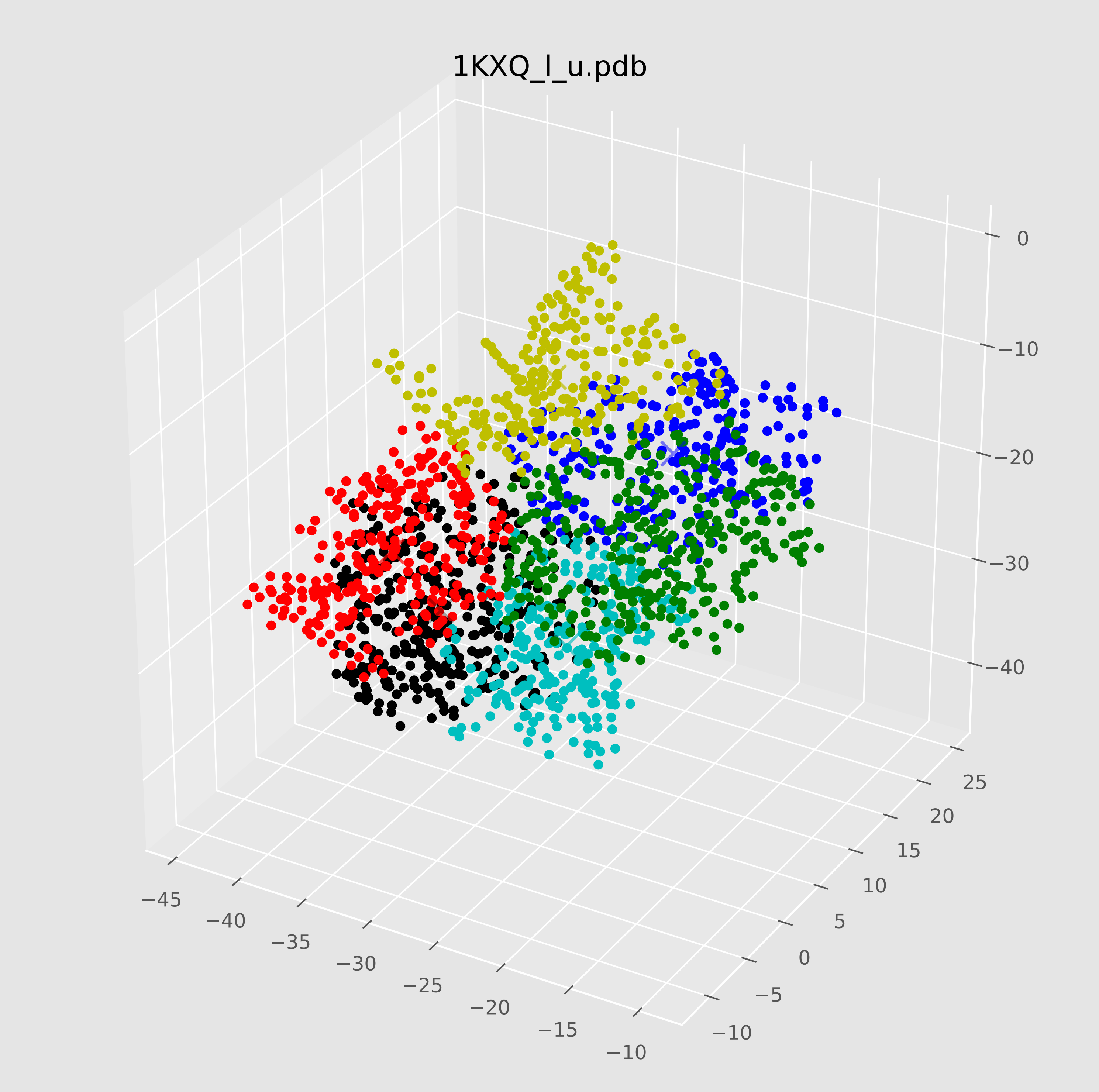

### 1KXQ_r_u.pdf

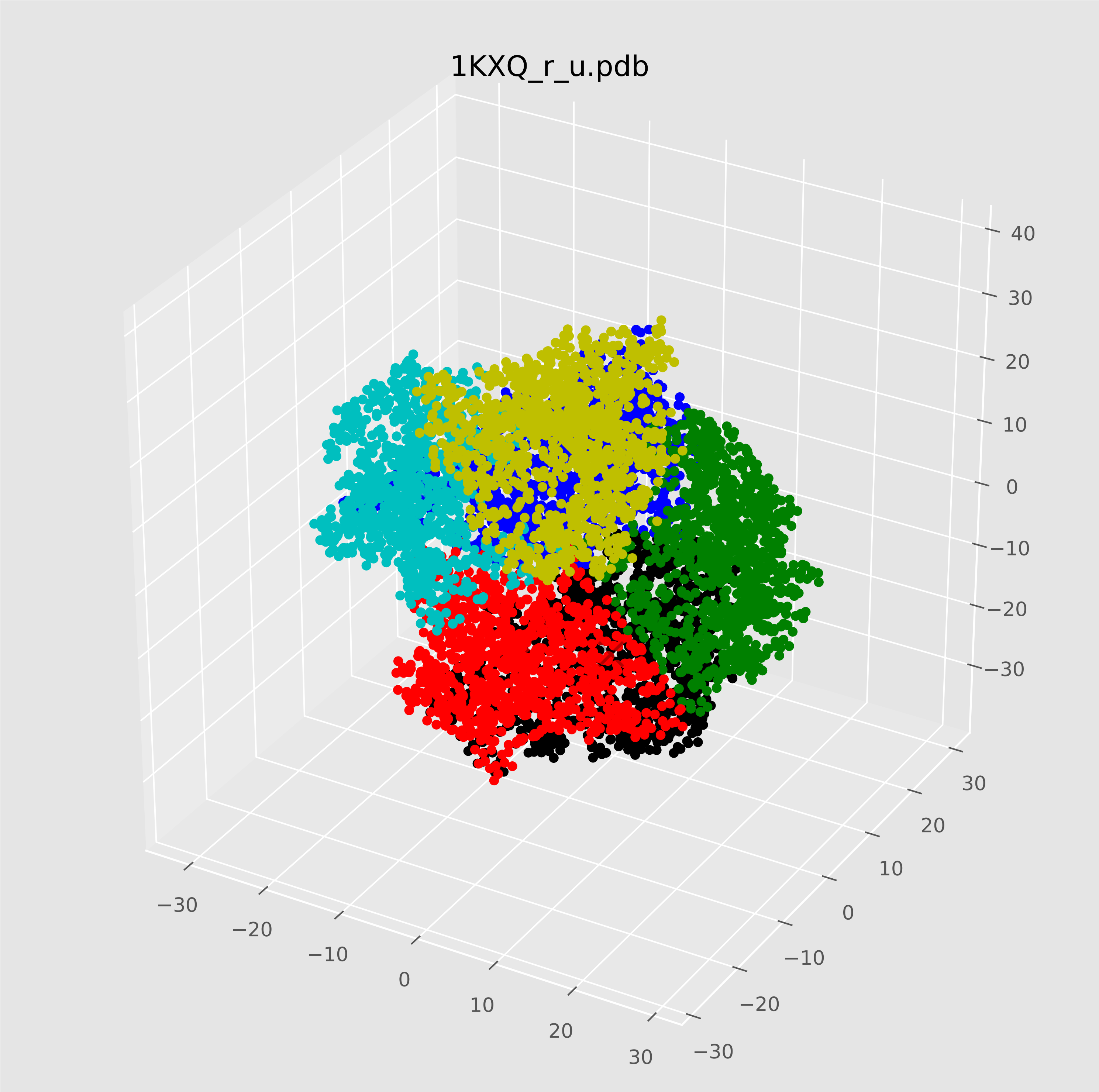

### 1M10_l_u.pdf

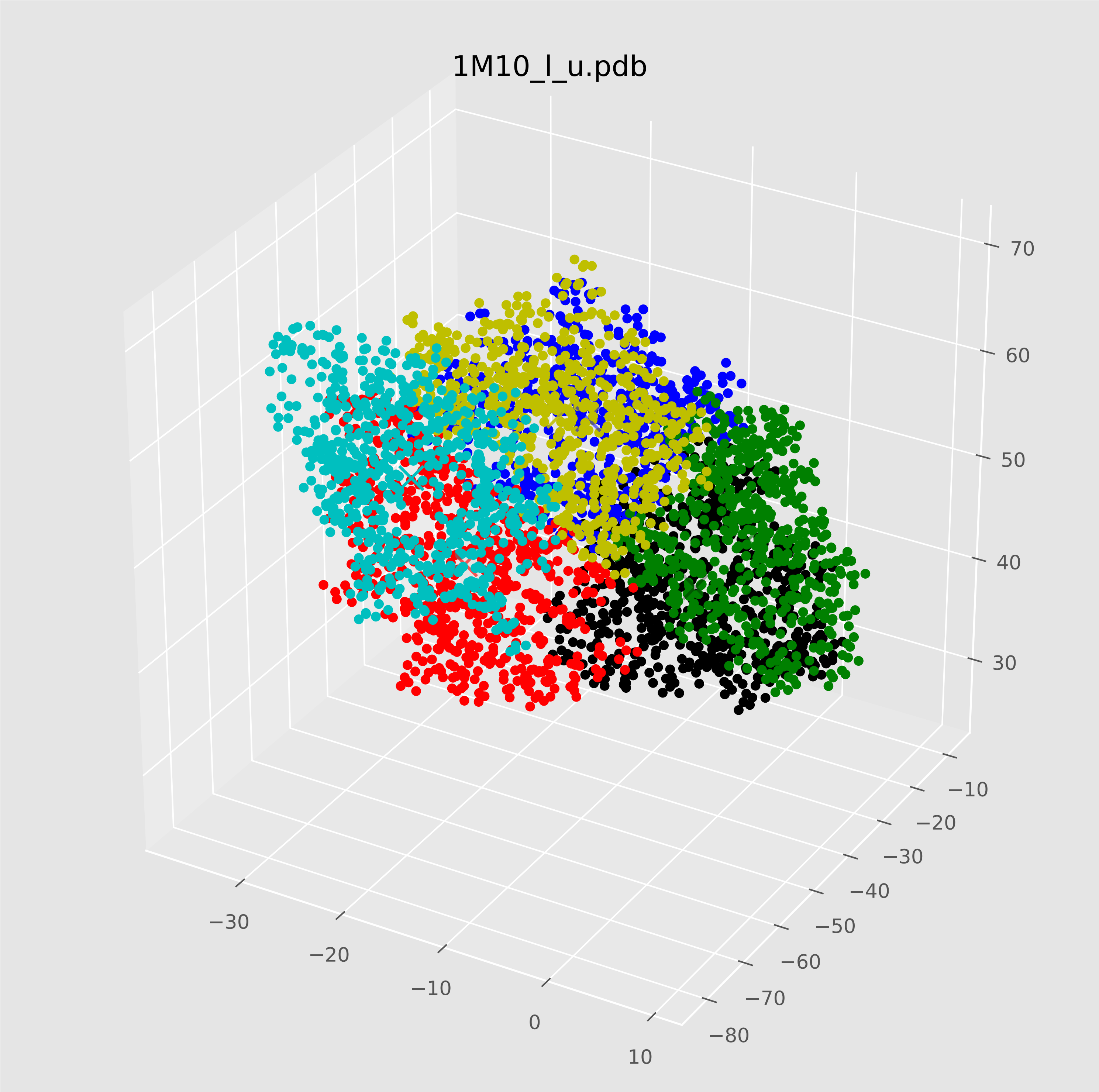

### 1M10_r_u.pdf

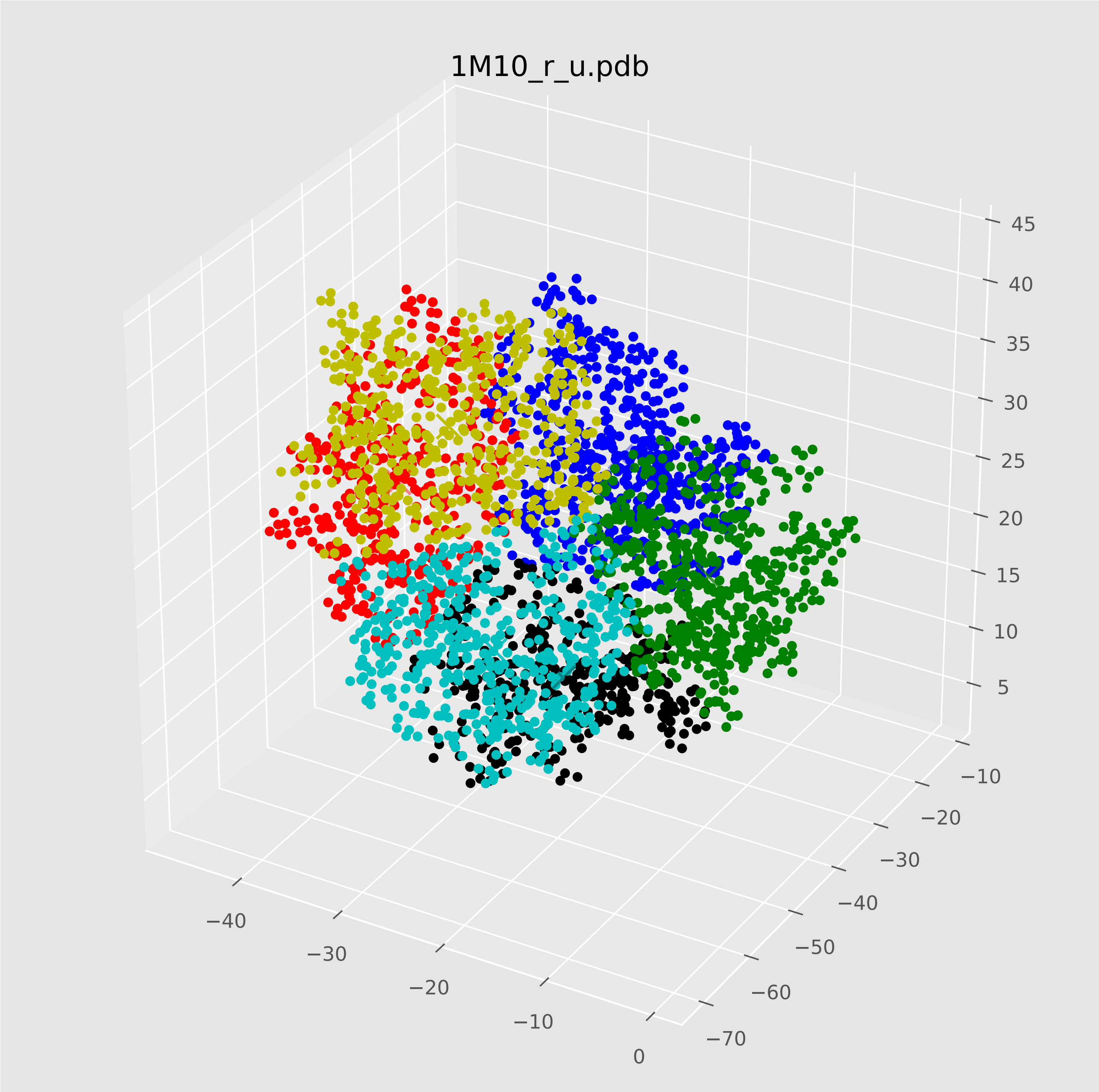

### 1MQ8_l_u.pdf

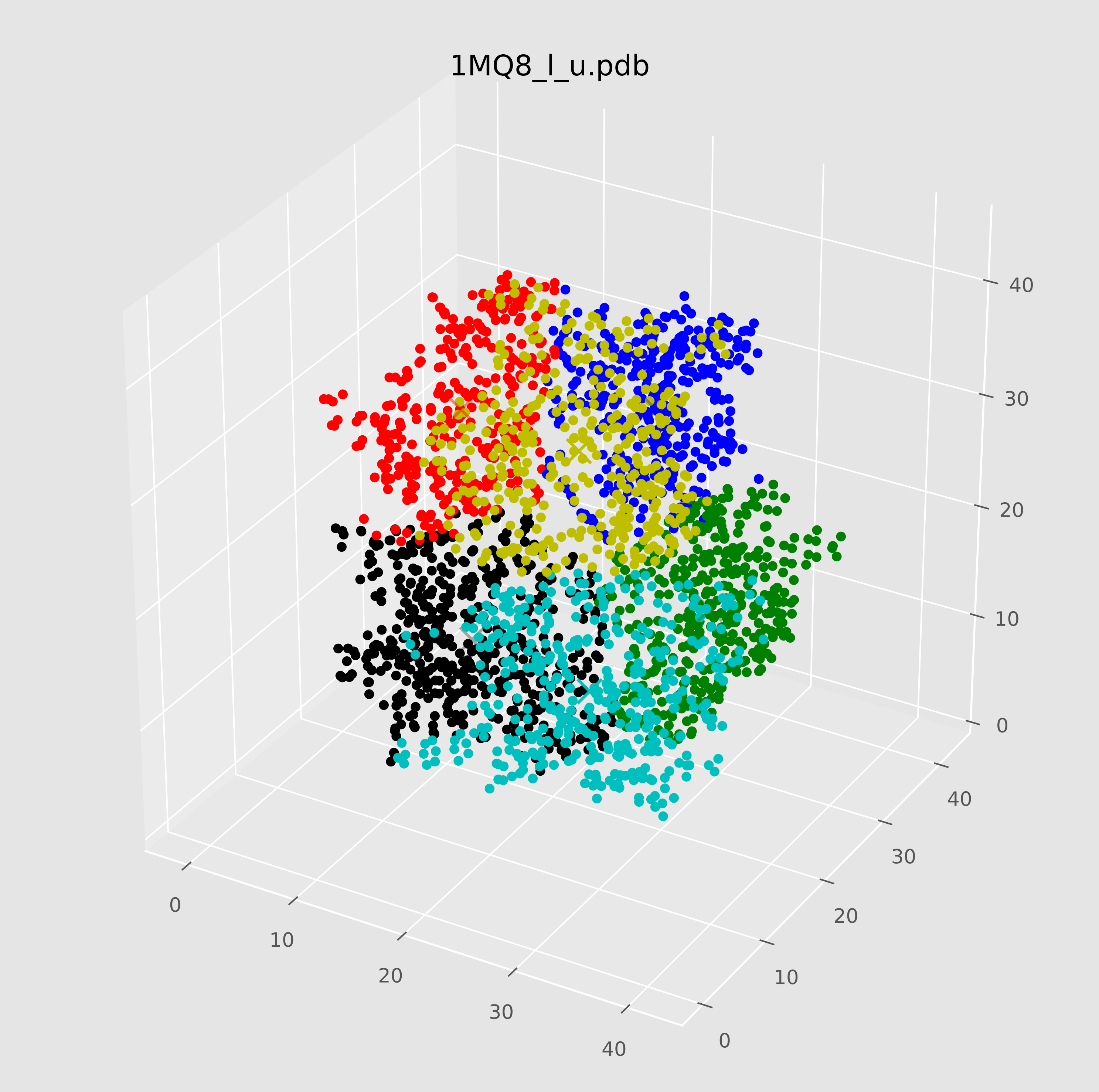

### 1MQ8_r_u.pdf

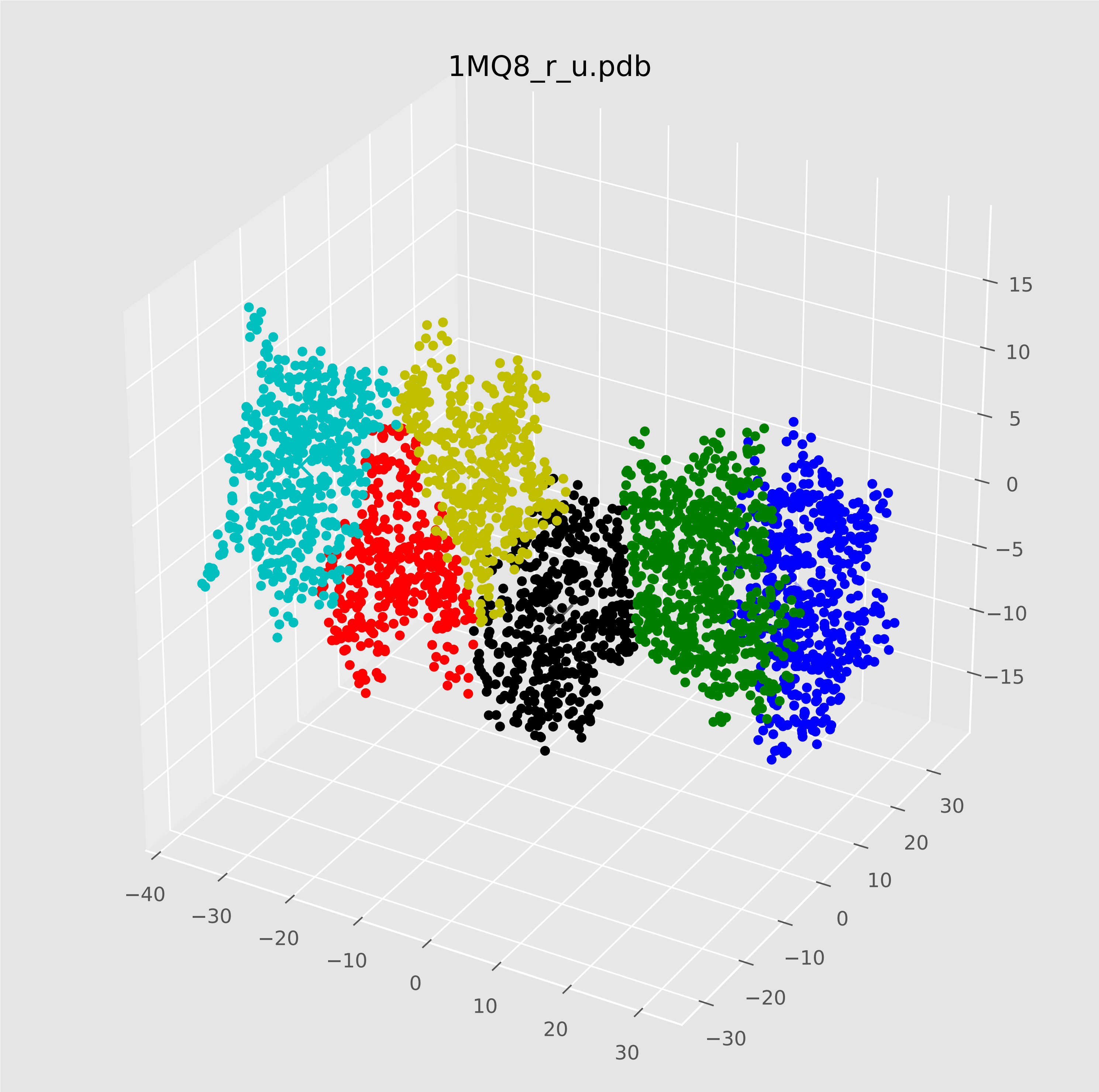

### 1T6B_l_u.pdf

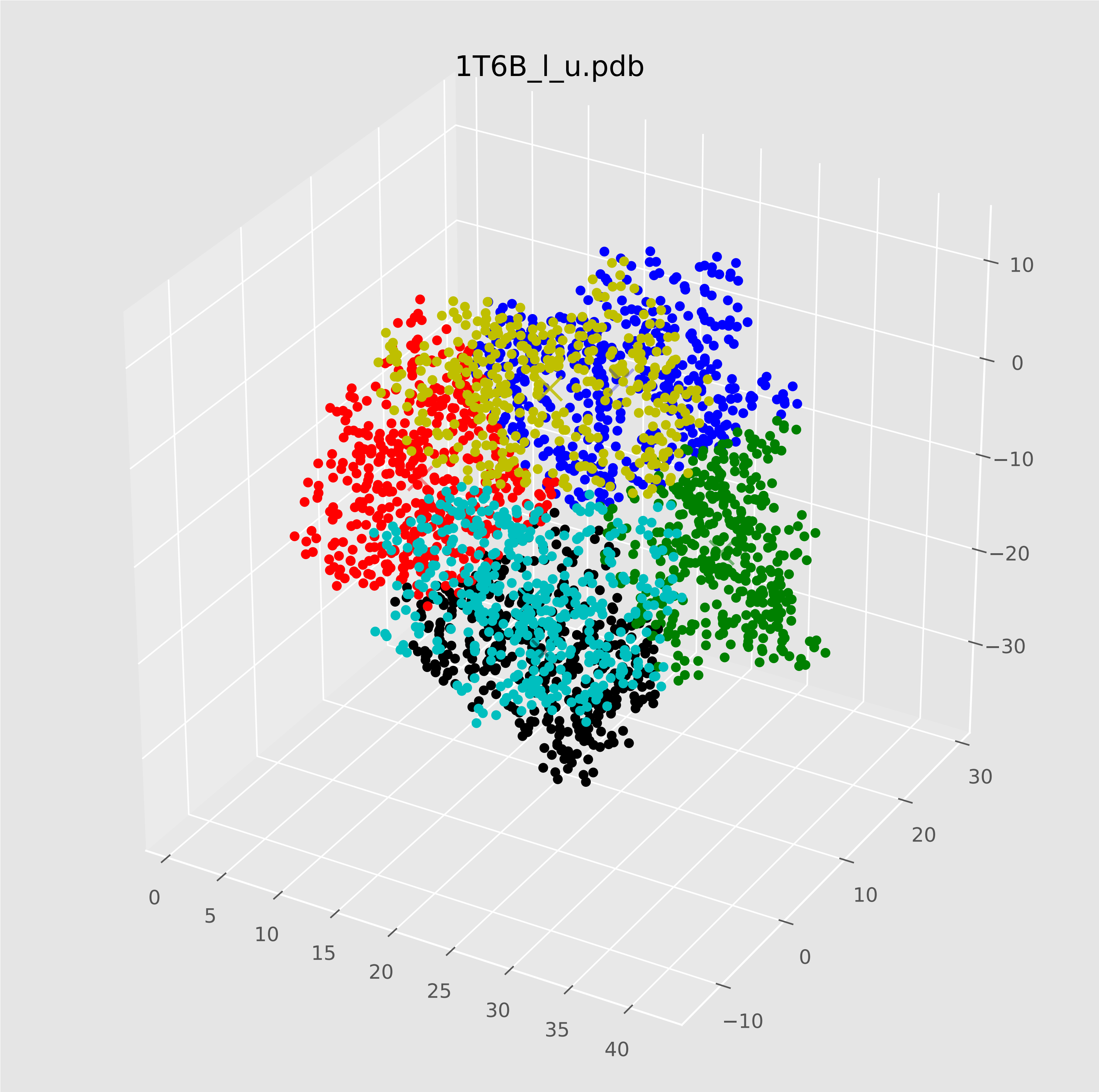

### 1T6B_r_u.pdf

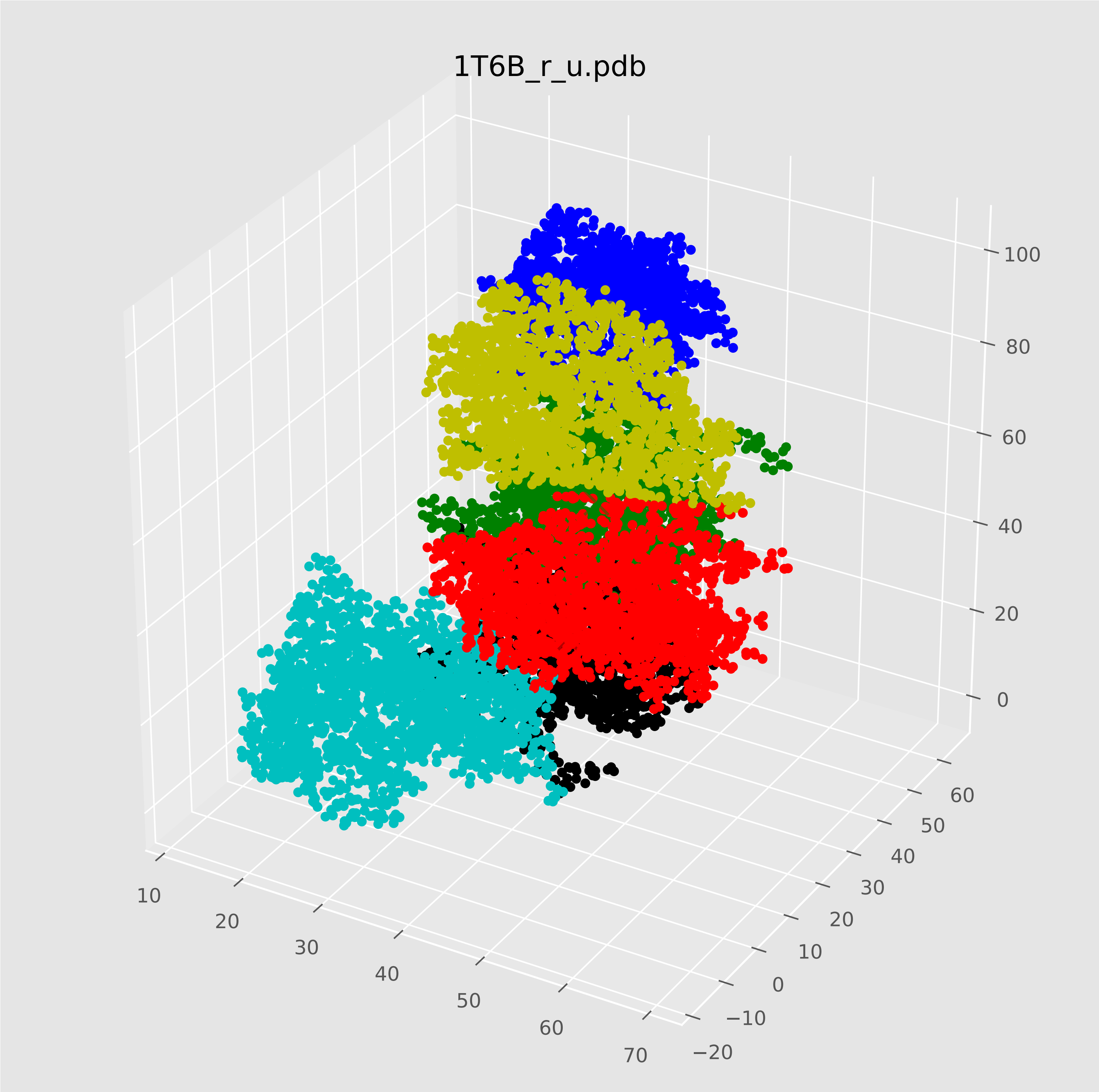

### 2FJU_l_u.pdf

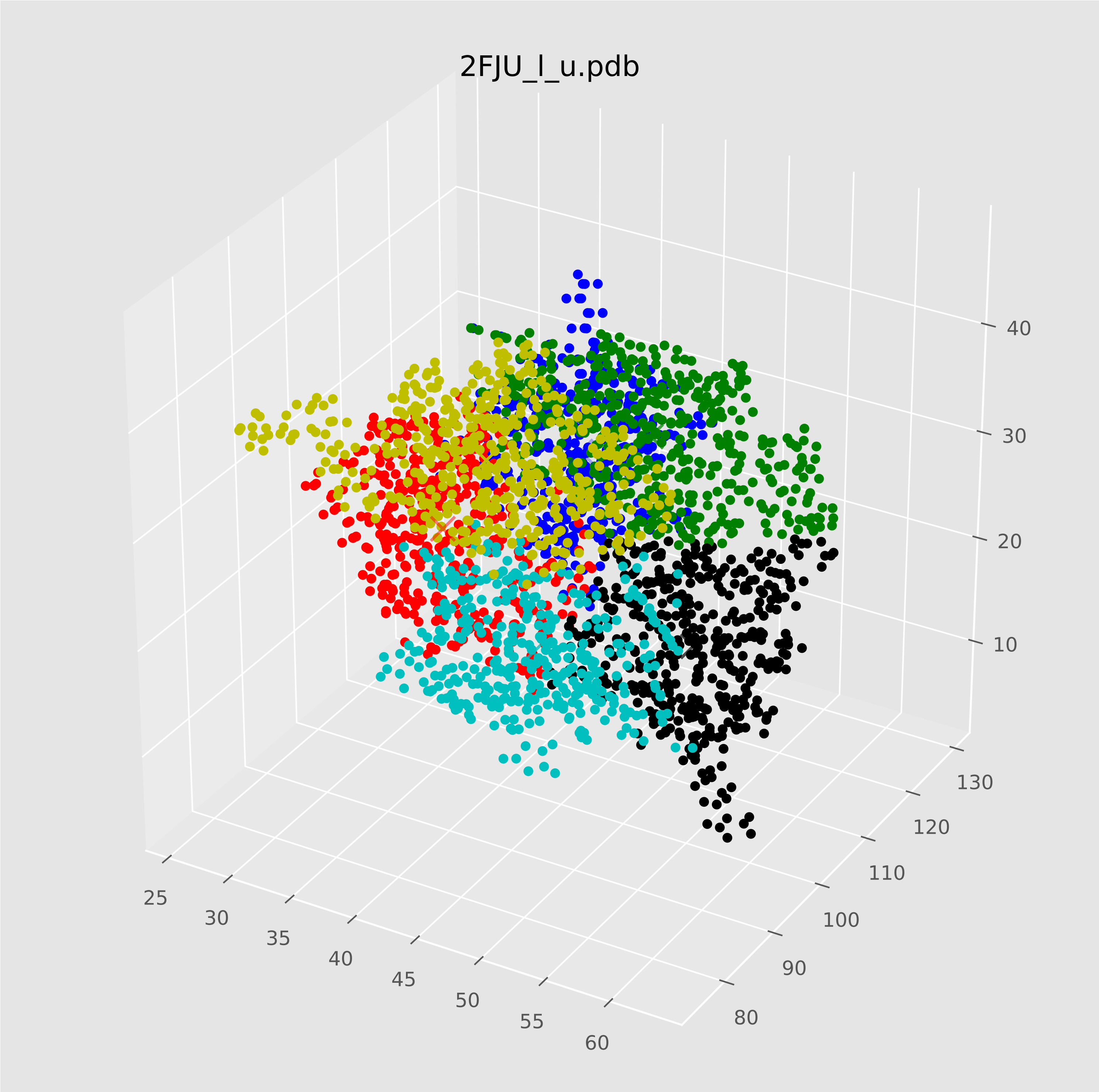

### 2FJU_r_u.pdf

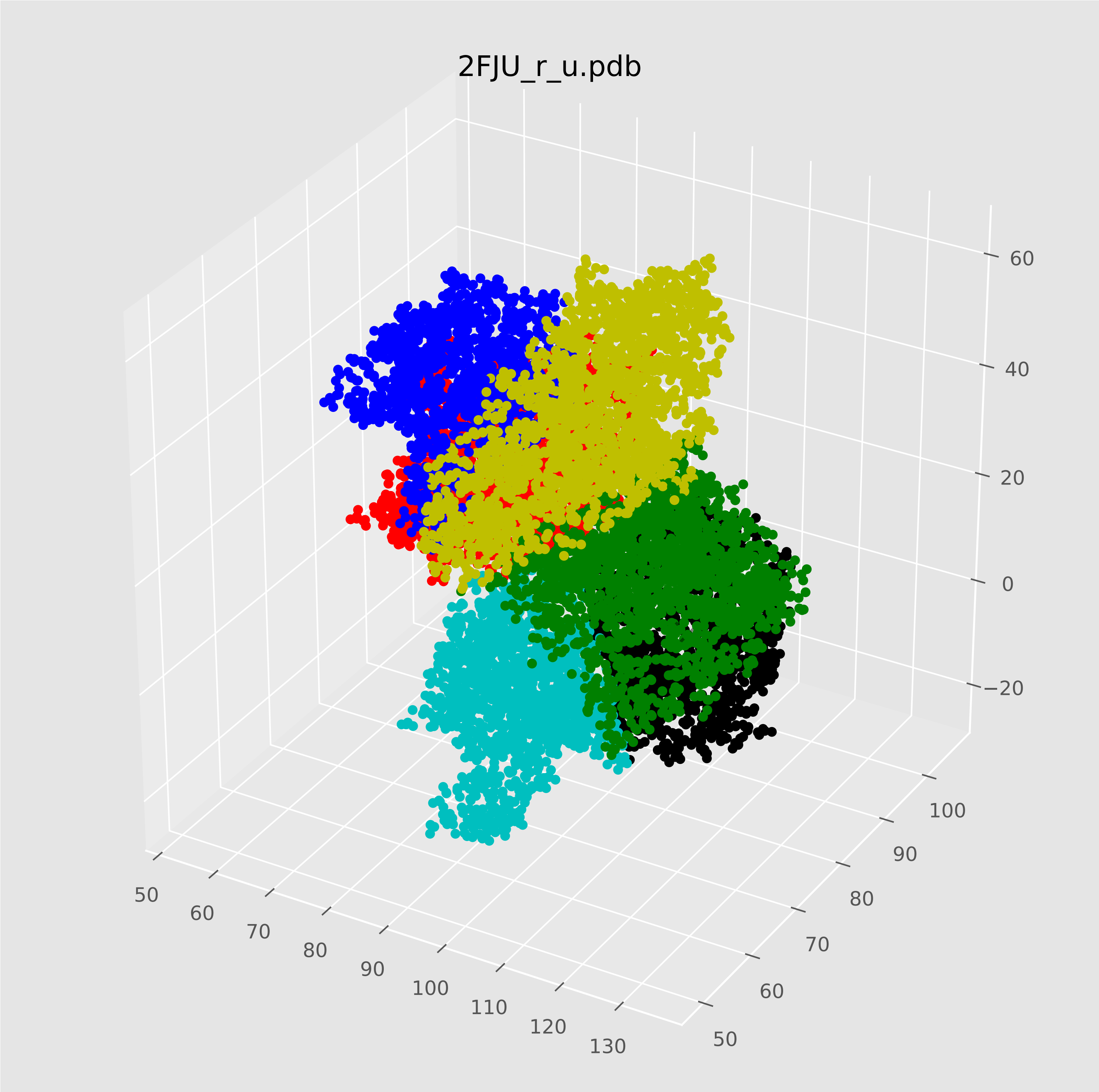

### 2HLE_l_u.pdf

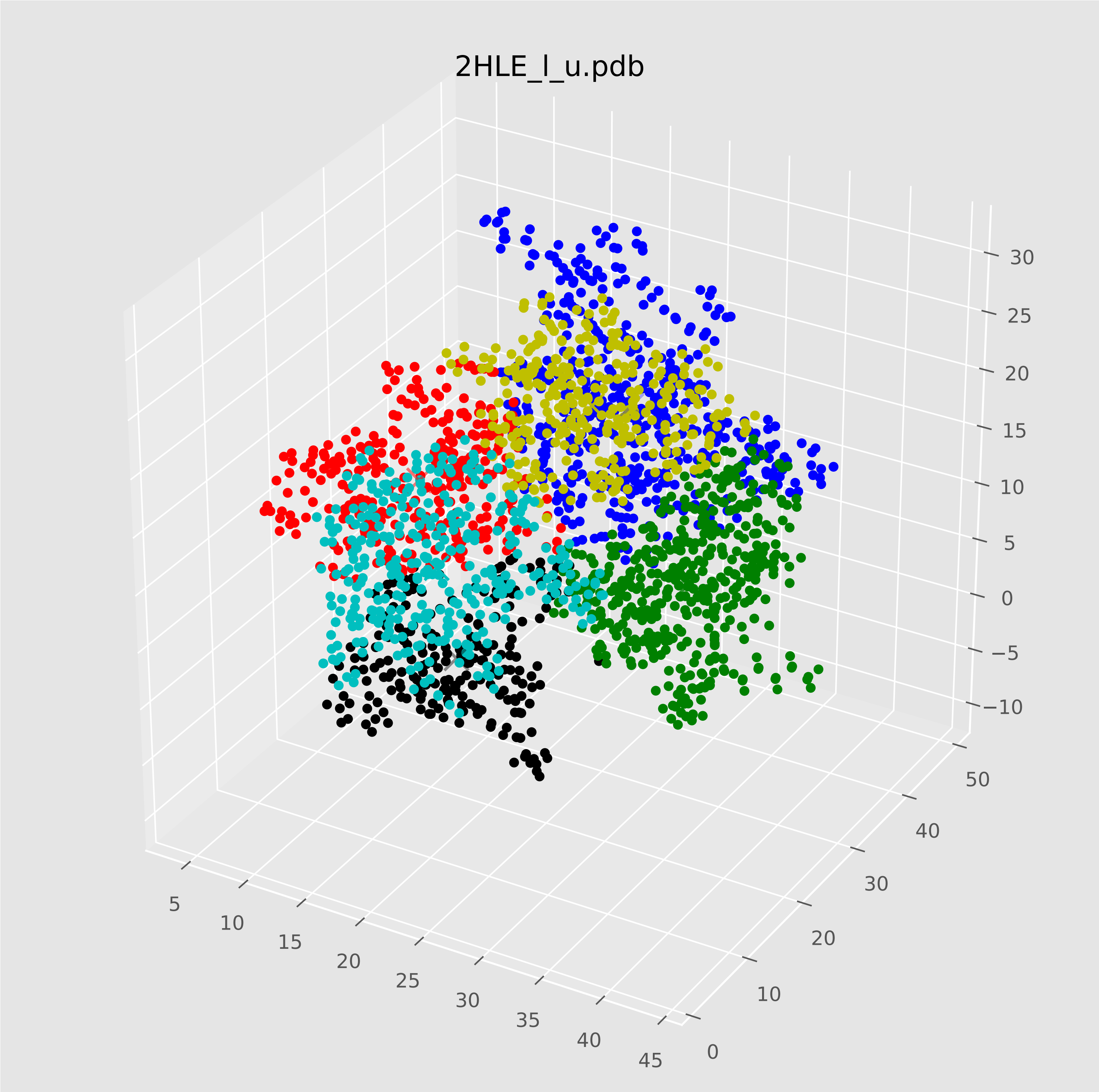

### 2HLE_r_u.pdf

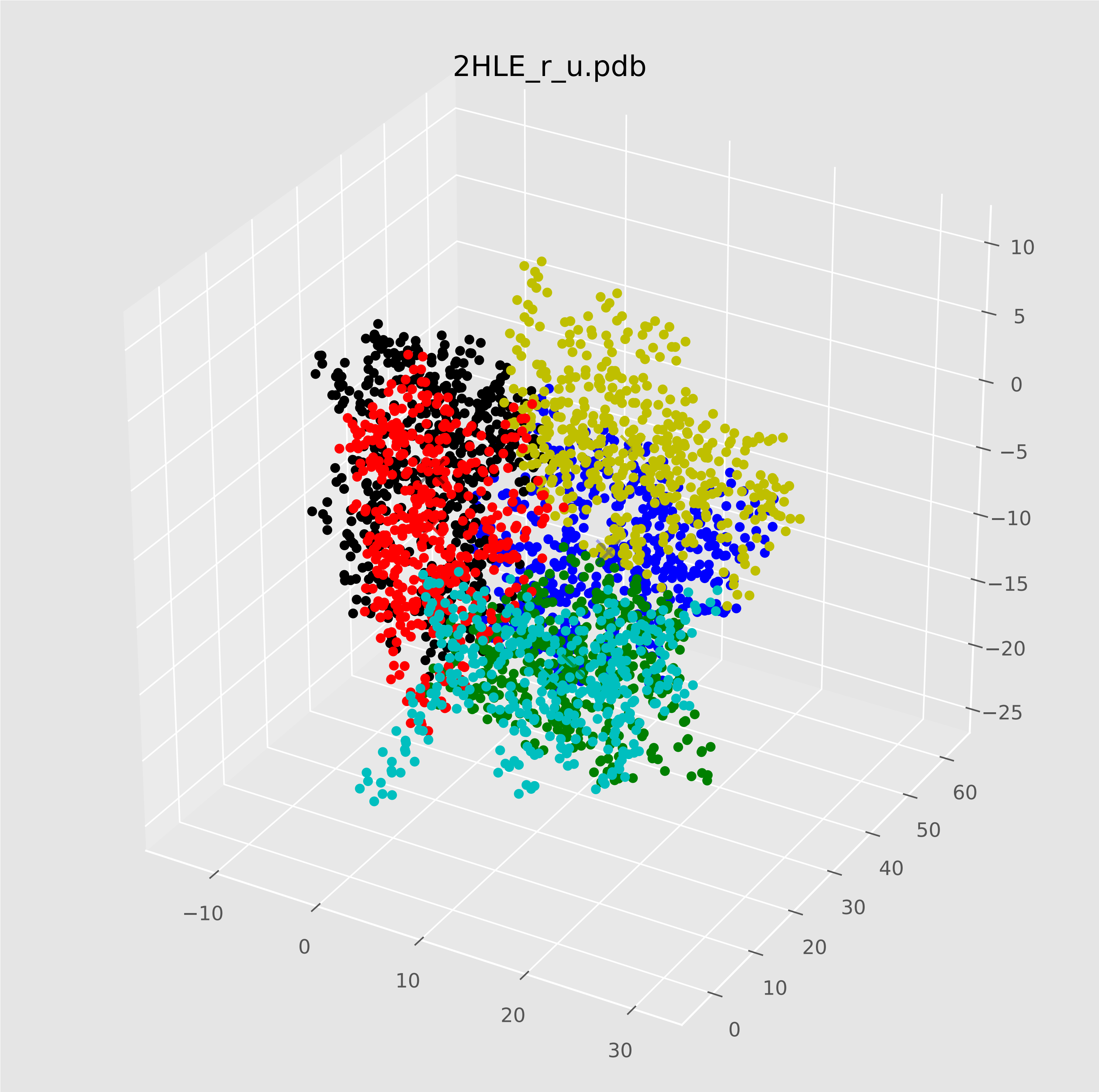

### 3CPH_l_u.pdf

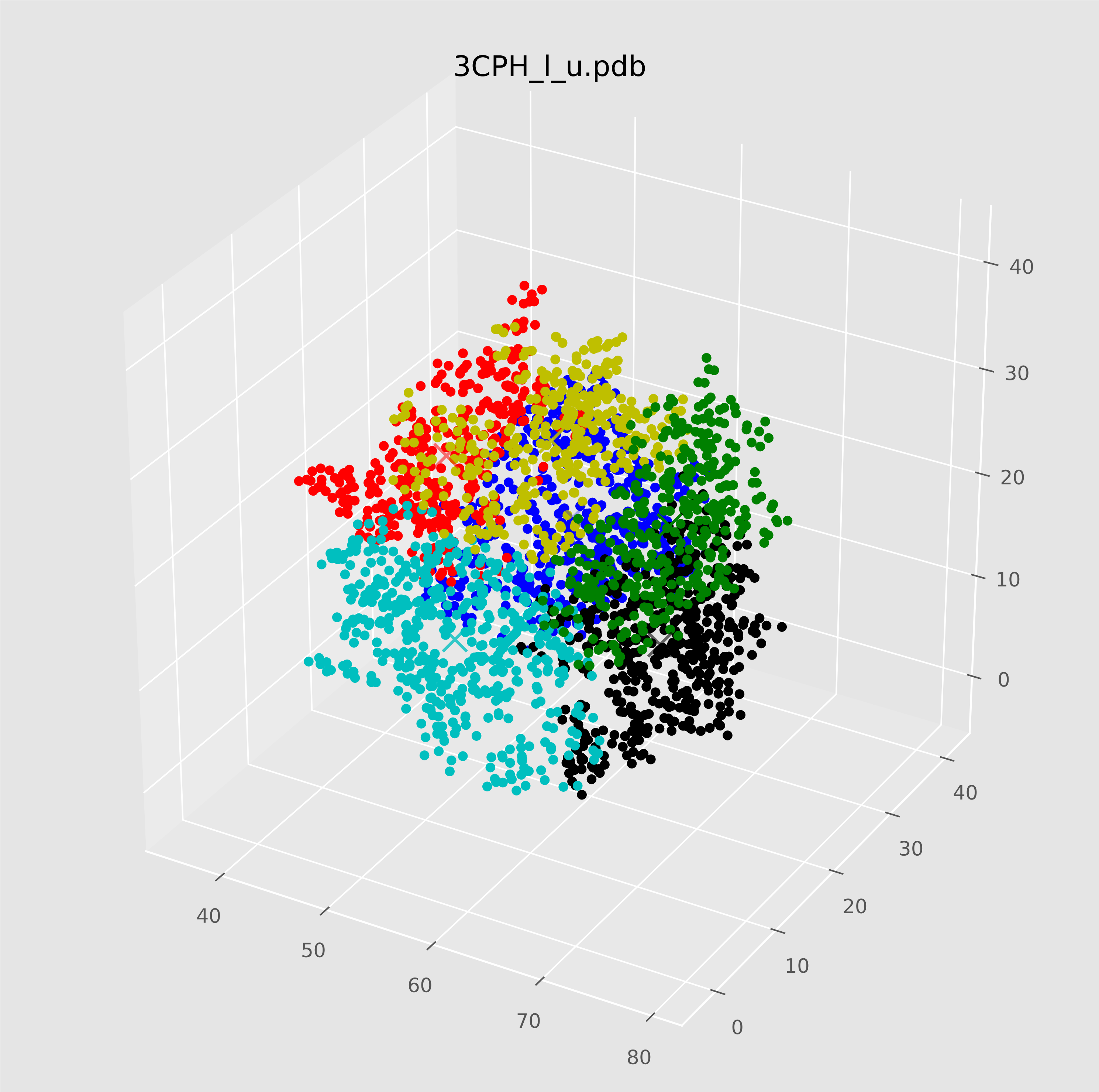

### 3CPH_r_u.pdf

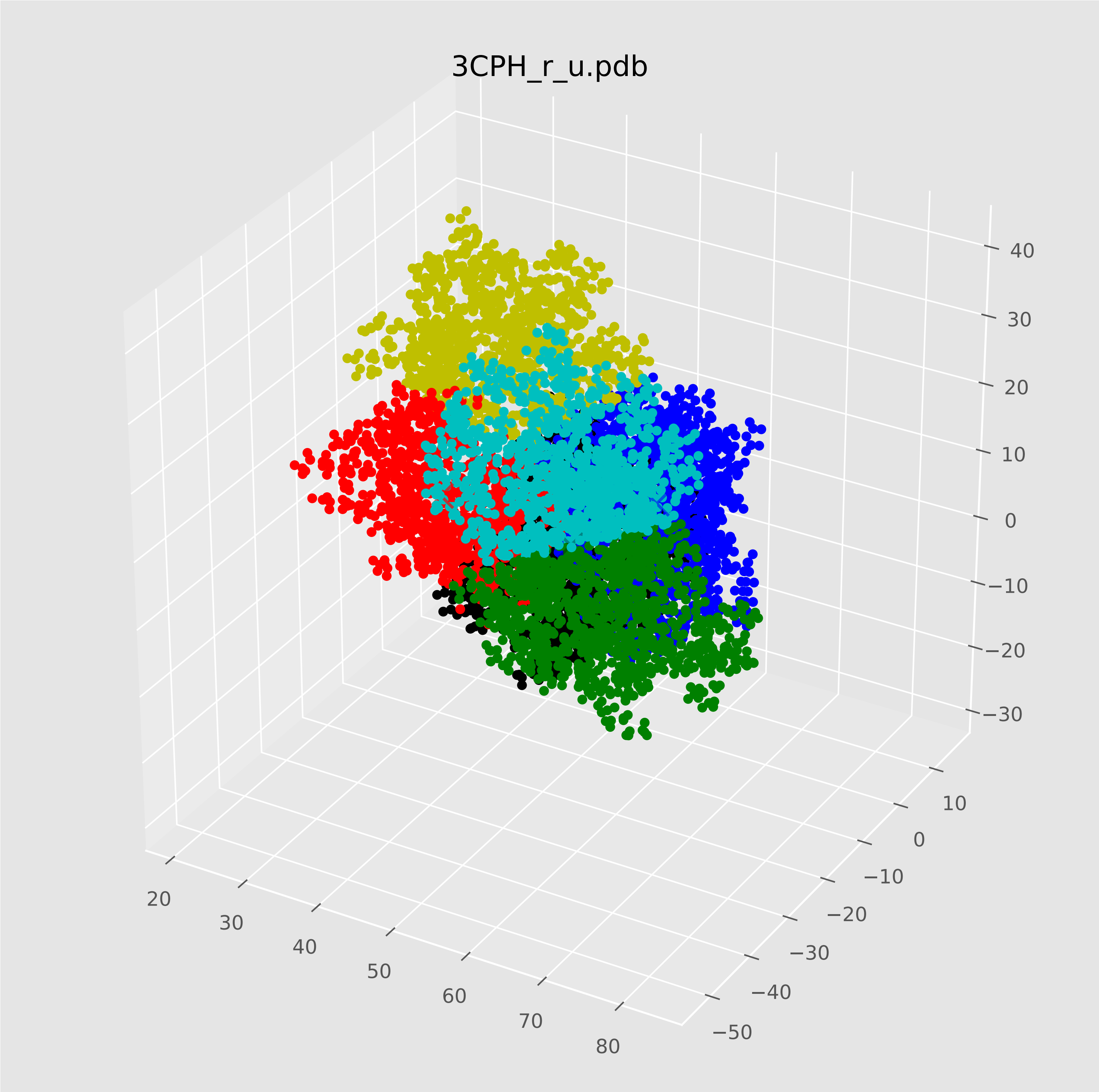
